# Chromatin information content landscapes inform transcription factor and DNA interactions

**DOI:** 10.1101/777532

**Authors:** Ricardo D’Oliveira Albanus, Yasuhiro Kyono, John Hensley, Arushi Varshney, Peter Orchard, Jacob O. Kitzman, Stephen C. J. Parker

**Affiliations:** Department of Computational Medicine & Bioinformatics, University of Michigan. Ann Arbor, USA; Department of Human Genetics, University of Michigan. Ann Arbor, USA; Tempus Labs, Inc. Chicago, IL, Chicago, USA

## Abstract

Interactions between transcription factors (TFs) and chromatin are fundamental to genome organization and regulation and, ultimately, cell state. Here, we use information theory to measure signatures of TF-chromatin interactions encoded in the patterns of the accessible genome, which we call chromatin information enrichment (CIE). We calculate CIE for hundreds of TF motifs across human tissues and identify two classes: low and high CIE. The 10-20% of TF motifs with high CIE associate with higher protein-DNA residence time, including different binding sites subclasses of the same TF, increased nucleosome phasing, specific protein domains, and the genetic control of both gene expression and chromatin accessibility. These results show that variations in the information content of chromatin architecture reflect functional biological variation, with implications for cell state dynamics and memory.

## Main text

Chromatin is the association between DNA, RNA, and diverse nuclear proteins, including nucleosomes. It enables the ~2-meter human genome to be packaged inside the nucleus while allowing active genes and their corresponding regulatory elements to remain accessible (*2*). Nucleosome positioning is an essential property of chromatin architecture and has been shown to have both passive and active roles in transcription factor (TF) binding (*3–5*). Understanding TF-chromatin interactions is therefore critical to dissect regulatory circuits that lead to differences in transcriptional activity across species, tissues, stimulatory, and genetic contexts. Information theory provides a powerful framework to quantify ordered patterns in data (*6*) and has been successfully used to characterize genome-wide DNA methylation patterns (*7*). Here, we hypothesized that local chromatin architecture encodes rich signatures of TF interactions and developed information-theoretical tools to measure these patterns in human tissues.

We first aimed to quantify patterns of chromatin accessibility around TF-chromatin interactions. We reasoned that TF binding creates a localized impact on chromatin architecture, which may result in TF-specific signatures. To measure chromatin architecture, we focused on the assay for transposase-accessible chromatin using sequencing (ATAC-seq) (*8*), that can simultaneously quantify both TF and nucleosome signatures, which are reflected in the ATAC-seq fragment length patterns. This chromatin architecture can be visualized using V-plots (*9*),which show the aggregate ATAC-seq fragment midpoints around TF binding sites and can result in a stereotyped “V” pattern of points for bound TFs with well-phased adjacent nucleosomes (Fig. 1A, upper plot). The extent of organization in the V-plot can be measured using Shannon’s entropy equations (*6*) to quantify information. We therefore calculated information content (*1*) of the ATAC-seq fragment size distribution around TF binding sites as a way to quantify V-plot organization (Fig. 1A, middle plot). To adjust for potential bias arising from non-uniform ATAC-seq fragment coverage across the V-plot, we devised a metric called chromatin information enrichment (CIE) (*1*) (Fig. 1A, middle and lower plots). We summarized CIE into a single value, named feature V-Plot Information Content Enrichment (f-VICE), which represents the CIE at landmark TF and nucleosomal positions across the V-plot (*1*), which are expected to have high CIE when the nucleosomes are phased around the TF binding site (Fig. 1A, lower plot). Therefore f-VICE quantifies the degree of chromatin architecture organization around a TF.

**Fig. 1.**
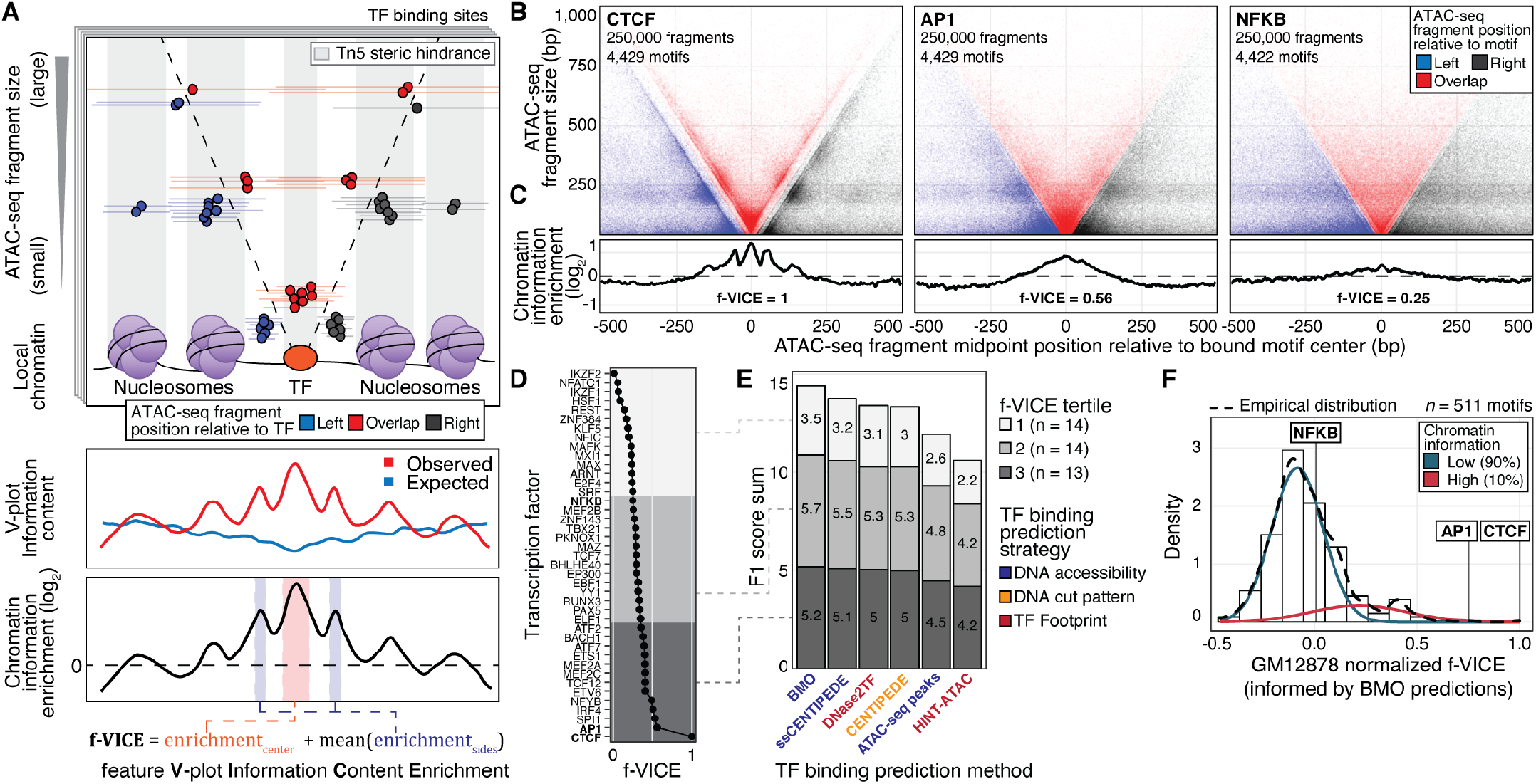
Information content of TF-chromatin interactions. (**A**) Upper: TF binding impacts the chromatin architecture and the observed ATAC-seq fragment distribution around TF binding sites. Middle and bottom: calculation of CIE and f-VICE. (**B-C**) V-plots and CIEs of CTCF, AP-1, and NFKB (GM12878 ATAC-seq data generated in this study). V-plots were downsampled to highlight differences in chromatin architecture (but not for f-VICE calculation). (**D**) f-VICEs calculated for TFs with GM12878 ChIP-seq data. (**E**) F1 score sum of TF binding prediction algorithms. (**F**) Normalized GM12878 BMO-informed f-VICE distribution.

We initially focused on the GM12878 lymphoblastoid cell line, for which there is high-quality, deeply-sequenced ATAC-seq data (*8*) and 41 TF chromatin immunoprecipitation followed by sequencing (ChIP-seq) experiments that pass our inclusion criteria (Table S1) (*1, 10*). To increase our ability to detect TF-chromatin interactions, we generated an independent GM12878 ATAC-seq dataset with higher signal-to-noise ratio (Fig. S1). Using these datasets, we created V-plots and calculated f-VICEs centered on bound motif instances for 41 TFs. The ATAC-seq fragment pattern was most ordered around CTCF, a known chromatin organizer (*11*), where we detected clusters of fragments distributed periodically in a “V” pattern indicating nucleosome phasing (Fig. 1B-C, S2). CTCF f-VICE was highest among the 41 TFs (Fig 1D). Other TFs, exemplified by AP-1 and NFKB, had diverse f-VICEs (Fig. 1B-D, S2). These patterns were consistent across independent ATAC-seq libraries, indicating the robustness of the f-VICE metric (Fig. S2). These results indicate extensive differences in TF-chromatin interactions, which are captured in the CIE patterns.

One alternative to determine f-VICEs for TFs without ChIP-seq data is to rely on binding predictions using chromatin accessibility data. This motivated us to first evaluate the performance of current TF binding prediction algorithms. Most algorithms search for footprints, which are regions of low chromatin accessibility embedded within larger accessible regions, thought to be caused by cleavage protection from bound TFs (*12–14*). However, a recent report indicated that ~80% of TFs do not have footprints (*15*). Hence, we developed BMO, an unsupervised method to predict TF binding using negative binomial models of chromatin accessibility (*16–18*) and co-occurring motifs (*19*), without relying on footprints (*1*). We benchmarked BMO and other methods (*12–14, 20*) using TF ChIP-seq data from GM12878 and HepG2 (Table S1). Overall, the footprint-agnostic methods (BMO, CENTIPEDE, and a custom implementation of CENTIPEDE, called ssCENTIPEDE (*1*, Supplementary Text) outperformed footprint-based methods on most (median of 81% across datasets) tested TFs, particularly on those with lower f-VICEs (Figs. 1E, S3-7; Supplementary Text). These findings indicate that TF binding is more accurately predicted using a simple chromatin accessibility model tuned to each TF motif.

Having determined that our BMO footprint-agnostic method is among the most accurate for predicting TF binding, we proceeded with predictions to estimate f-VICEs for TFs without ChIP-seq data. BMO-predicted f-VICEs were significantly correlated with f-VICEs calculated from TF ChIP-seq data across all datasets (Pearson’s ρ≥0.72, *p*≤1e-10; Fig. S8). We therefore concluded that BMO can be used to estimate f-VICEs without ChIP-seq data and performed BMO TF binding predictions to calculate f-VICEs for 540 non-redundant (*1*) TF motifs (Tables S2-3). We used high-quality ATAC-seq datasets from four additional human tissues (pancreatic islets (*21*), pancreatic islet sorted alpha and beta cells (*22*), and CD4+ cells (*23*); Table S1), selected by applying a strategy that uses the highly stereotyped chromatin architecture in ubiquitous and conserved CTCF/cohesin binding sites to measure sample quality (Fig. S9) (*1*). We normalized f-VICEs within each sample (*1*) to control for differences in bound motif predictions and overall chromatin accessibility (Fig. S10). Among the 540 motifs, we observed a mixture of two f-VICE distributions and therefore used a mixture of two gaussians to fit the data (*1*). The median percentage of motifs associated with high f-VICEs across datasets was 14% (Figs. 1G, S11), which is comparable to the percentage of motifs associated with DNase footprint protection across datasets (median=19%) from another study (*15*) and supports our conclusion that footprint-based algorithms will not perform well on the majority (median of 86% across datasets) of TFs. Together, these results reinforce the use of footprint-agnostic methods like BMO for accurately calculating f-VICE.

TF residence time, which corresponds to the duration of DNA binding for a TF, is an important biophysical measurement that can influence TF activity (*3, 24*). Based on the high f-VICEs for CTCF and AP-1 and low f-VICE for NFKB (Fig. 1C-D), which agree with the known residence times for these TFs (Table S4), we hypothesized that CIE correlates with residence time. We correlated BMO-informed f-VICEs with previously measured fluorescence recovery after photobleaching (FRAP) data from mammalian cell lines (Table S4), which provide an upper bound of TF residence time (*25, 26*). Using a robust linear regression to protect against outlier influence, we found that f-VICE was significantly associated with FRAP recovery times in all samples (β≥0.7, Bonferroni adjusted *p*≤0.001; Figs. 2A, S12). This suggests that TFs associated with high CIE have longer residence times.

**Fig. 2.**
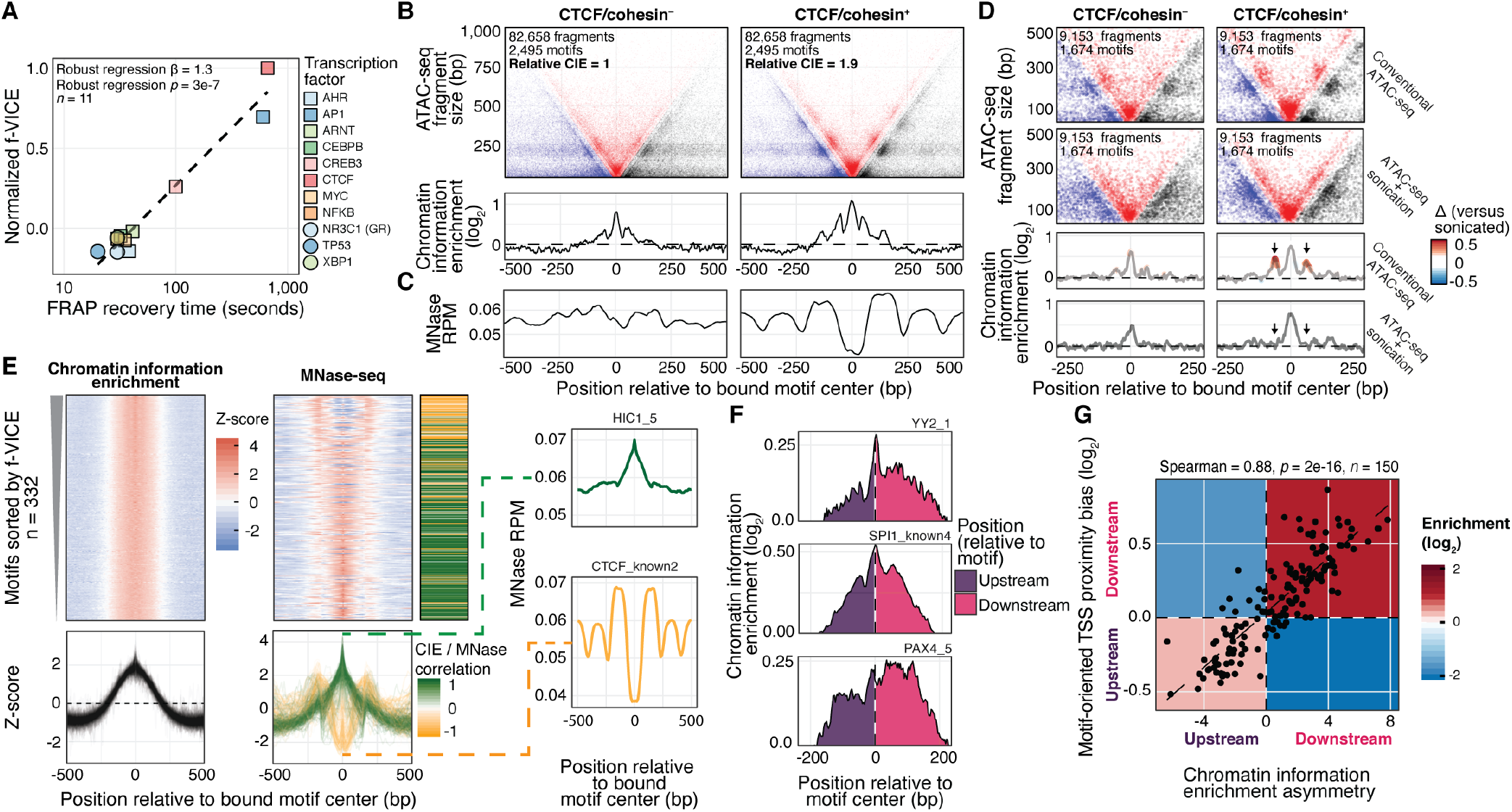
Chromatin information informs residence times and TF-nucleosome interactions. (**A**) Correlation of FRAP recovery times and GM12878 f-VICEs. Dashed line, linear model fit. (**B**) V-plots and CIEs of CTCF/cohesin^+^ and CTCF/cohesin^−^ motifs. (**C**) GM19238 MNase-seq reads per million mapped reads at the same motifs. (**D**) CTCF/cohesin^+^ and CTCF/cohesin^−^ motifs in the sonicated and conventional GM12878 ATAC-seq data. Colors, differences relative to sonicated. (**E**) Left: CIE and MNase-seq profiles (*k*-means cluster three). Middle: Heatmap of MNase and CIE Z-score correlations. Right: Example motifs with positive and negative CIE/MNase correlation. (**F**) Top 3 motifs with CIE asymmetry Z-scores in GM12878. (**G**) Scatter plot of motif-oriented TSS position bias and CIE asymmetry in TSS-proximal motifs. Enrichments calculated by permuting the signs of observed values (*n*=10,000).

A recent study found that cohesin has a residence time 10-to 20-fold higher than CTCF (*26*). We reasoned this difference could be reflected in the local chromatin architecture and calculated the CIE of the GM12878 lymphoblastoid cell line CTCF binding sites with and without the presence of cohesin (CTCF/cohesin^+^ and CTCF/cohesin^−^), controlling for potential confounding biases (Fig. S13A) (*1*). CTCF/cohesin^+^ had 1.9-fold higher CIE compared to CTCF/cohesin^−^ (Figs. 2B, S13B), indicating these distinct CTCF occupancy classes have different CIE signatures. We next compared the nucleosome positioning signals inferred from lymphoblastoid cell line micrococcal nuclease sequencing (MNase-seq) profiles (Table S1). Only the CTCF/cohesin^+^ class had phased nucleosomes around the binding site (Figs. 2C, S13C), consistent with longer residence times associating with nucleosome phasing. To experimentally validate these results, we generated chromatin accessibility data using a modified ATAC-seq protocol with an additional sonication step (*1*) to disrupt the fragment size information (Fig. S14). There were no detectable nucleosome phasing patterns in the motif-flanking CIE signature of the sonicated sample (Fig. 2D; see vertical arrows). These results are complementary to our residence time results above in that they show our CIE approach can capture differences even when subsets of a single TF have different residence times.

To systematically characterize the association between CIE and nucleosome positioning, we compared GM12878 CIE patterns across TF motifs to lymphoblastoid MNase-seq profiles. First, we used *k*-means clustering (*1*) to divide motifs into broad CIE categories. We found three clusters representing a continuum of CIE at the motif region (Fig. S15A). Clusters one and two had lower CIE at the motif compared to motif-flanking regions, while cluster three had the highest CIE at the motif (Fig. 2E, S15A-B) and encompassed >95% of the high f-VICE motifs (Fig. S15C-D). Notably, we observed two distinctly anti-correlated MNase-seq signal patterns for the motifs in cluster three (Fig. 2E). These two distinct patterns are consistent with TFs binding at the center of the nucleosome dyad or between phased nucleosomes (*5, 27*). CIE and MNase signals were anti-correlated at high f-VICE motifs (Fig 2E; yellow-green heatmap), indicating that the highest CIE TFs associate with nucleosome phasing. We quantified nucleosome phasing (*1*) and found that it was significantly correlated with f-VICE in clusters two and three (Spearman’s ρ≥0.42, *p*≤1e-7; Fig. S16). These results suggest that TF-chromatin interaction patterns are driven by TF residence time, resulting in distinct CIE signatures.

Previous reports suggested that a subset of TFs directionally bind DNA, with potential effects on gene regulation (*12, 28, 29*). To investigate this further, we extended our information content analyses to quantify CIE asymmetry (*1*). Of the 540 motifs tested, 150 had significantly asymmetric CIE (Bonferroni corrected *p*<0.05; Figs. 2F, S17A). The direction of CIE asymmetry was significantly correlated with the direction of the nearest TSS relative to each motif instance (Spearman’s ρ=0.66, *p*=2e-16; Fig. S17B). To determine if asymmetric CIE was an artifact of TSS proximity, we calculated CIE asymmetry separately for TSS-proximal (≤1 kb) and TSS-distal (≥10 kb) motif instances. The TSS-distal and TSS-proximal CIE asymmetry directions agreed significantly more than expected by chance (111/150, binomial test *p*=4e-9; Fig. S17C-D), suggesting that CIE asymmetry is intrinsic to the TF motif. The magnitude of asymmetry was higher in TSS-proximal motifs (Fig. S17D), suggesting that TSS proximity amplifies TF CIE asymmetry. Accordingly, the correlation between nearest TSS direction and CIE asymmetry was stronger at TSS-proximal motifs (Spearman’s ρ=0.88, *p*=2e-16; Fig. 2G). These results indicate that directional binding is an intrinsic property of TF-chromatin interactions.

We next aimed to investigate cross-tissue differences in CIEs. We performed an unsupervised hierarchical clustering of motif f-VICEs and found that it recapitulated the expected tissue grouping (Fig. 3A). A recent study demonstrated that NF-KB (p65) residence time is determined by its DNA-binding domain (DBD) (*30*), which motivated us to ask if DBDs are associated with CIE. We assigned DBDs and protein domains to motifs and designed a permutation-based rank test to calculate domain f-VICE enrichments (*1*). We observed both common and tissue-specific f-VICE enrichments, including IRF and ETS in blood-related samples, PAX in islet-related samples, and HMG/SOX and FOX domains in HepG2 (FDR < 10%; Figs. 3B, S18). Our findings show the landscape of TF-chromatin interactions varies across tissues and reflects protein domain-level TF properties.

**Fig. 3.**
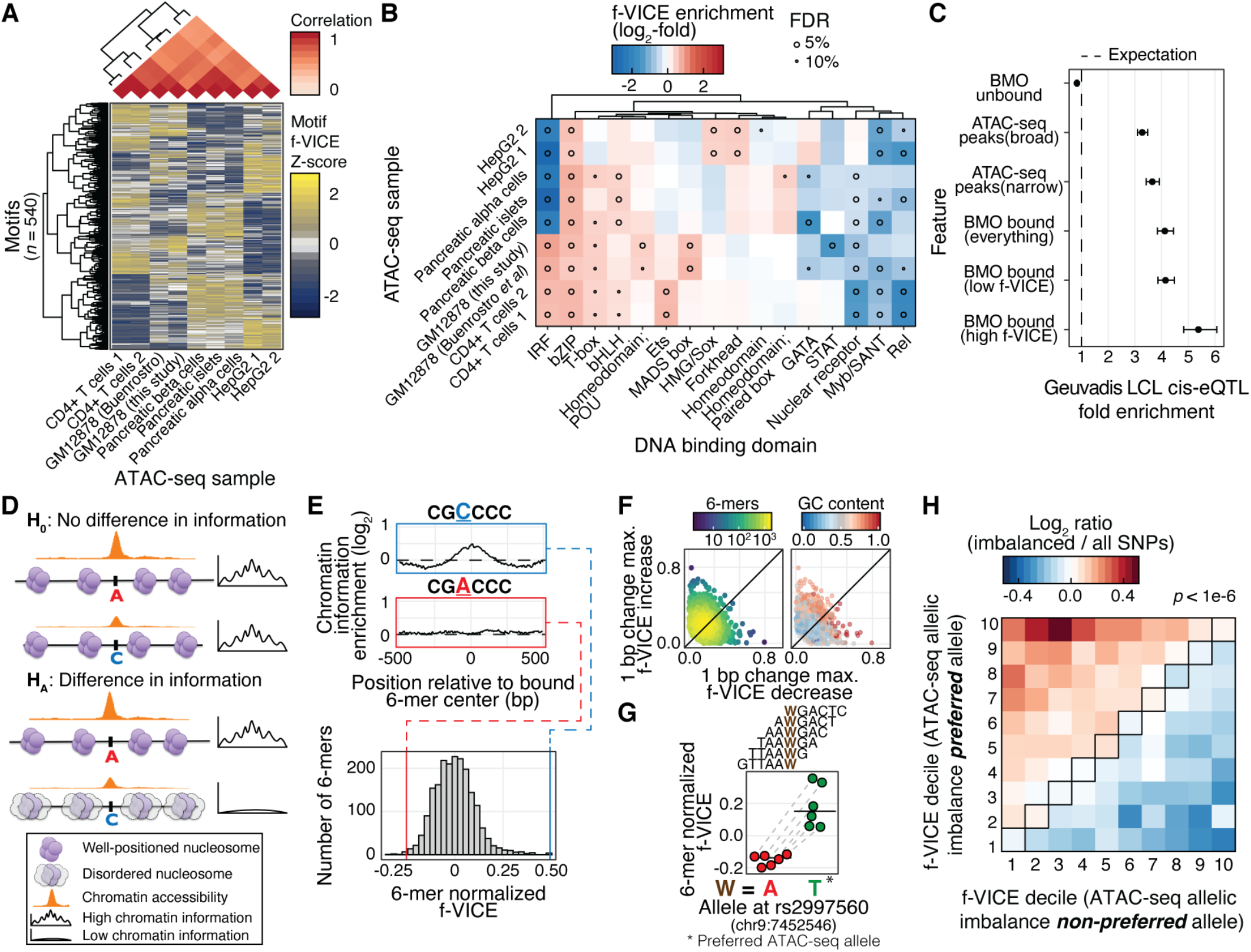
The chromatin information landscape of human tissues. (**A**) Hierarchical clustering of f-VICE Z-scores. (**B**) f-VICE enrichments across DBDs. (**C**) LCL cis-eQTLs enrichments. Error bars, effect size SD. (**D**) Hypothesis schematic. (**E**) Upper: Two 6-mers with 1bp difference in sequence. Lower: pancreatic islets 6-mer normalized f-VICE distribution. (**F**) Range of f-VICE differences associated with 1-bp difference in 6-mer sequence. (**G**) Predicted f-VICE change associated with rs2997560 in pancreatic islets. Horizontal bars, median. (**H**) Log_2_ ratio of f-VICE decile changes associated with the preferred and non-preferred alleles of imbalanced SNPs versus all tested SNPs in pancreatic islets.

The prevalence of tissue-specific differences in CIEs led us to examine the role of high f-VICE TFs in regulating gene expression. We calculated the enrichment of the motifs categorized as high or low f-VICE in GM12878 (Fig. 1F) to overlap lymphoblastoid cis-expression quantitative trait loci (cis-eQTLs) datasets (*31, 32*), which represent gene expression genetic control regions. High f-VICE motifs had 15-30% higher (median=24%) fold-enrichment in cis-eQTLs compared to low f-VICE motifs (Figs. 3C, S19A), but no differences in eQTL effect sizes (Fig. S19B). These results indicate that high f-VICE TFs are more likely to mediate genetic effects on gene expression, but not their magnitude.

Given that high f-VICE TFs have highly ordered chromatin (Fig. 1), high predicted residence times (Fig. 2A, B, S12), and nucleosome phasing properties (Fig. 2E, S16), we hypothesized that their regulatory effects (Fig. 3C) could result from acting as or recruiting pioneer factors that induce chromatin accessibility (*12, 33*). If true, we would expect increased CIE for single nucleotide polymorphism (SNP) alleles with increased chromatin accessibility (*i.e.* with ATAC-seq allelic imbalance; Fig. 3D). We performed a motif-agnostic approach to calculate the f-VICEs associated with every DNA 6-mer, controlling for differences in chromatin accessibility (*1*). This strategy allows the interrogation of genetic variants by determining the DNA 6-mers formed by each allele and their corresponding f-VICEs. DNA 6-mers have a distribution of f-VICEs (Figs. 3E; S20A) and GC-pure 6-mers had the highest f-VICEs (Fig. S20B), which is consistent with GC-rich sequences driving enhancer activity (*34*) and suggest that high GC-content regions represent anchors of nuclear architecture. Notably, a single basepair change can lead to large differences in 6-mer f-VICEs (Fig. 3E-F, S20C-E), suggesting that genetic variation impacts CIE. To test this, we determined f-VICEs for 6-mers formed by both alleles at SNPs with significant ATAC-seq allelic imbalance (binomial test *p*<0.05) in GM12878 and pancreatic islets (*1*). The preferred ATAC-seq alleles were significantly biased to form higher f-VICE 6-mers compared to the less favored allele in all samples (permutation test *p*<3e-4; Fig. 3G-H, S21). These findings support a model where TFs with potential pioneer-like properties bookmark regions of the genome to allow binding of other migrant-like TFs (*12, 33*). Notably, TF motifs that are predictive of binding without any chromatin accessibility data (based solely on the motif match score) have significantly higher f-VICEs in GM12878 and HepG2 (robust linear regression *p*≤0.001; Fig. S22). This suggests that high f-VICE TFs, particularly CTCF, are more likely to bind any strong motif regardless of its underlying accessibility, while the remaining TFs require motifs located in already accessible regions. Collectively, our results show that application of information theory methods to chromatin profiles captures a dynamic landscape of TF-chromatin interactions, with implications for cell state memory and gene regulation.

## Acknowledgments

We thank members of the Parker Lab, L. J. Scott, P. Freddolino, P. Wittkopp, M. Burmeister, G. Higgins, P. Pereira, M. Puthenveedu, and J. Brancho for helpful comments. Sequencing was performed at the UM Sequencing Core Facility.

## Author contributions

ROA: Analyzed data, designed computational experiments, wrote the manuscript. YK: Generated ATAC-seq datasets. JH: Implemented computational algorithms. AV: analyzed eQTL data. PO: calculated ATAC-seq allelic imbalance. JK: designed ATAC-seq experiments. SCJP: designed experiments, analyzed data, wrote the manuscript, and supervised all aspects of the project.

## Data and materials availability

Code and scripts are available (github.com/ParkerLab/chromatin_information). ATAC-seq data is deposited in GEO (GSE135074).

