## Supplementary materials for "Chromatin information content landscapes inform transcription factor and DNA interactions"

**This PDF file includes:**

Materials and Methods
Supplementary Text
Figs. S1 to S22
Table S4

**Other Supplementary Materials for this manuscript include the following:**

Tables S1 to S3

### Materials and Methods

GM12878 cell culture

We cultured GM12878 cells following the ENCODE GM12878 cell culture protocol (encodeproject.org/documents/1bb75b62-ac29-4368-9855-68d410e1963a), with added plasmocin (Invivogen, San Diego, CA; 50 ug/mL) to the growth media to prevent mycoplasma contamination.

GM12878 ATAC-seq data generation

We conducted ATAC-seq as described in (*35*) using a home-made Tn5 that we synthesized as described in (*36*). For each replicate we incubated 250,000 cells with 12.5 uL of 1:1 mix of Tn5 enzyme that carry 5-methylC-MEDS-A oligos and MEDS-B oligos at 37° C for 30 minutes in a 50 uL reaction. We column-purified the tagmented DNA using the Zymo DNA Clean & Concentrator-5 kit (Zymo Research, Irvine, CA) and constructed Illumina sequencing library using the Kitzman lab custom indexing primers. We PCR-amplified a total of 11 cycles until amplification curve reached its mid-log phase (⅓ to ½ of max signal), and then purified the PCR products using SPRI beads prepared as in (*37*) and eluted in 22 uL of TTE8 buffer. Sequencing was performed on an Illumina HiSeq 4000 platform at the University of Michigan Sequencing Core and a total of ~33 million paired-end 52 bp reads were generated.

Sonicated GM12878 ATAC-seq data generation

For each replicate we incubated 250,000 cells with three different concentrations of enzyme (0.2X, 1X, and 5X; 1X corresponds to 2.5 uL of Tn5 that carry 5-methylC-MEDS-A oligos) at 37° C for 30 minutes in a 50 uL reaction. We column-purified the tagmented DNA using the Zymo DNA Clean & Concentrator-5 kit (Zymo Research, Irvine, CA), and sonicated to ~350 bp using the Covaris M220 sonicator (peak incident power - 50W; duty factor - 20%; cycles per burst - 200; treatment time - 60 sec). We constructed Illumina sequencing library using the ACCEL-NGS Methyl-seq DNA Library kit (Swift Biosciences #DL-ILMMS-12; revision 160106) with the following modifications to the manufacturer's protocol: 1) We skipped "Ligation" and "Post-ligation SPRI" steps (pg. 10-11), as the 5' end of the fragments had already been tagged during the transposition step. Accordingly, we eluted DNA with 20 uL of TTE8 (10 mM Tris-HCl, 0.1 mM EDTA, 0.05% Tween-20, pH 8) for Post-Extension SPRI step (pg. 10), instead of 15 uL, to adjust for the difference in volume before proceeding to the "Indexing PCR" step (pg. 11); 2) We used 2:1 beads:sample ratio for "Post-Extension SPRI" step (pg. 10) and 1.8:1 beads:sample ratio for "Post-PCR SPRI" step (pg. 12); and 3) For indexing PCR, we used the Kitzman lab custom primers (barcode plate #5) to prime the P5 end and the "IndexD7XX" primers (Swift Biosciences #DI-ILMMS-48) to prime the P7 end. We PCR-amplified a total of 14 cycles for 0.2X, 1X samples and 16 cycles for 5X samples until amplification curve reached its mid-log phase (⅓ to ½ of max signal), and then purified the PCR products using SPRI beads prepared as in (*37*) and eluted in 22 uL of TTE8 buffer. Sequencing was performed on an Illumina HiSeq 2500 platform at the University of Michigan Sequencing Core and a total of ~33 million paired-end 126 bp reads were generated.

ATAC-seq data processing

Reads were trimmed for barcodes and aligned to the hg19 reference human genome using BWA mem (v. 0.7.15) (*38*) similarly to our previous study (*39*), with additional parameters *-I 200,200,5000* to avoid larger ATAC-seq fragments being discarded. We removed duplicate alignments using Picard (broadinstitute.github.io/picard) and retained properly paired and uniquely mapped alignments with high mapping quality using samtools view (v. 1.3.1) (*40*) with flags *-f 3 -F 4 -F 8 -F 256 -F 1024 -F 2048 -q 30*. We called broad and narrow peaks using MACS2 (v. 2.1.1.20160309) (*41*) with flags *-g hs --nomodel --shift -100 --extsize 200 -B [--broad] --keep-dup all* and kept peaks that did not intersect blacklisted regions by the ENCODE consortium due to poor mappability (sites.google.com/site/anshulkundaje/projects/blacklists), using bedtools (v2.26.0) (*42*), and that reached 5% FDR. All data was processed uniformly using Snakemake (*43*).

Motif processing

We used the PWM scans from (*39*). Briefly, we used biallelic SNPs and short indels from the 1,000 Genomes project (release v5) (*44*) to generate comprehensive scans with FIMO (*45*), using the background nucleotide frequencies from hg19 and a $p$<1e-4. We only kept motif instances that intersected mappable regions and did not intersect blacklisted regions. In order to reduce motif redundancy, we performed PWM clustering in our motif database using the *matrix-clustering* tool from RSAT (*46*), with parameters *-lth cor 0.7 -lth Ncor 0.7*. For each of the 540 clusters obtained, we retained the motif with the highest total PWM information content for downstream analyses.

V-plots, chromatin information enrichments, and f-VICEs

V-plots (*9*) were generated by creating a matrix of aggregated fragments from the selected set of genomic features (motifs), removing all instances that overlap within ±500 bp of each other. We used the script *measure_signal* (using flags *-r 500*) , which is part of a suite of tools to analyze ATAC-seq data we developed for this study (github.com/ParkerLab/atactk). Each cell in the V-plot matrix outputted by *measure_signal* correspond to the number of fragment midpoints at a position relative to the feature center ($x$-axis) and fragment size ($y$-axis). We binned the matrix in the $x$-axis using a sliding window of 10 bp width, with 2-bp overlap, and summed all the values within the window corresponding to a given fragment size. For each $x$-axis bin, we calculated the normalized information content (I) of the corresponding fragment size distribution ($y$-axis) using the formula $I(x) = 1 - \frac{H(x)}{H_{max}}$, where $H(x)$ is the maximum-likelihood Shannon's entropy function implemented of the entropy R package (*47*), and $H_{max}$ is the Shannon's entropy of the information length (*i.e.* the range of fragment size distribution of the entire V-plot). To calculate the expected normalized information content, we randomly permuted the position and size labels in the fragments and repeated the steps outlined above.

Chromatin information enrichment was calculated as the $\log_2$ of the observed normalized information content divided by the expected normalized information content. For Fig. 1C, V-plots were downsampled to equal number of ATAC-seq fragments and motifs between TFs by selecting the top $n$ motifs ranked by the number of ATAC-seq fragments within ±500 bp, and then further downsampling to 250,000 fragments, where $n$ represents the smallest number of bound motifs among the plotted TFs. These downsampled V-plots were only used for visualization purposes and not used for the f-VICE calculations described below.

To obtain the feature V-plot information content enrichment (f-VICE) for each motif, we summed of the average chromatin information enrichment in the V-plot regions corresponding to the center (-25 to 25 bp) and proximal (-70 to -50 and 50 to 70 bp) chromatin information enrichment peaks referent to the CTCF V-plot, which correspond to small fragments spanning the TF binding site and to those positioned between the TF and the first pair of proximal nucleosomes, respectively (Fig. 1A). This value was then normalized across all motifs using the residuals of the linear model *f-VICE* $\sim \log_{10}(m) + \log_{10}(f)$, where *m* corresponds to the number of predicted bound motif instances for each motif and *f* corresponds to the total number of ATAC-seq fragments at the predicted bound motif instances for each motif (Fig. S10). The residuals for each sample were divided by the corresponding CTCF value in that sample to normalize it to 1. The linear model normalization was not performed in the ChIP-seq f-VICEs reported in Figs. 1D and S8 due to lack of data points to accurately fit the linear model.

Additional ATAC-seq samples selection

In addition to the GM12878 ATAC-seq dataset generated for this study, we analyzed an additional eight publicly available datasets corresponding to pancreatic islets (*21*, *22*), CD4+ cells (*23*), GM12878 (*8*), and HepG2 (*48*). With the exception of HepG2, these datasets were selected from a survey of all the public ATAC-seq datasets available until the end of 2017. We selected for our analyses datasets with at least 20 million high-quality autosomal reads and transcription start site (TSS) enrichment ≥ 6. In addition, we only retained samples with the stereotypical chromatin information enrichment indicative of nucleosome phasing at ubiquitous and conserved CTCF-cohesin binding sites. These regions provide a reference V-plot, with expected high accessibility and periodical chromatin information enrichment patterns in any high-quality sample. The ubiquitous and conserved CTCF-cohesin sites were defined as CTCF motifs overlapping ENCODE CTCF and Rad21 ChIP-seq peaks in at least in at least 54/59 (CTCF) and two (Rad21) different human tissues, located in bi-directionally mappable regions between human and mouse using bnMapper (*49*) that also corresponded to CTCF motif matches in the mm9 reference genome. To quantify samples, we defined our high-quality GM12878 dataset as reference and calculated the chromatin information enrichment correlation ≤ 200 bp from the motif center. Samples with correlation < 0.8 (Spearman) were discarded (Fig. S9). Finally, we only retained tissues/cells that had at least two samples that passed our stringent selection criteria. A list of all dataset accessions used in this study can be found in Table S2.

BMO transcription factor binding prediction

BMO builds on previous reports that the degree of chromatin accessibility around a motif (*16*–*18*) and the presence of co-occurring motifs (*19*) positively correlate with TF occupancy and uses per-TF negative binomial models of these two signals to estimate the likelihood of a TF motif instance being bound. Using all genomic matches for a given motif PWM, we calculated the number of ATAC-seq fragments overlapping a region ±100bp from the motif match, ignoring fragments that integrated within the motif coordinates. The latter step was performed to mitigate ATAC-seq bias, given that the nucleotide sequence in the motif regions is relatively constant across features and is more subject to assay-specific bias compared to the motif-flanking regions. We randomly selected 10,000 motif matches occurring outside ATAC-seq peaks to fit a background ATAC-seq fragment negative binomial distribution. We repeated this step 100 times and calculated the average mean and overdispersion. This approach is 1-2 orders of magnitude faster compared to fitting the negative binomial distribution on the entire set of motif matches outside ATAC-seq peaks and yielded identical results in a test using a representative subset of our data that accounted for the number of motif matches per PWM. We calculated the *p* values for the number of ATAC-seq fragments in all motif matches based on the fitted per-TF background distributions. Next, for each motif instance, we determined the number of co-occurring matches for the same PWM within ±100 bp and fitted a second negative binomial distribution on the co-occurring motifs. We combined the nominal *p* values of the ATAC-seq fragments and co-occurring motifs distributions by summing their Z scores (*50*) and corrected for multiple testing correction using the Benjamini-Yekutieli correction procedure (*51*). Motif instances were considered bound where the adjusted *p* value < 0.05.

CENTIPEDE

For each PWM scan result, we generated a strand-specific (relative to the motif orientation) base-pair resolution matrix encoding the number of Tn5 transposase integration events in a region ±100 bp from each motif occurrence using *make_cut_matrix* with parameters *-d -r 100*. This matrix and the motif PWM score were used as input for CENTIPEDE (v. 1.2), and a motif occurrence was considered bound if the outputted posterior probability was higher than 0.99. To calculate AUC-PRs, we used the posteriors outputted by the software as scoring metric. We developed *make_cut_matrix* as part of atactk (github.com/ParkerLab/atactk).

Signal-sum CENTIPEDE (ssCENTIPEDE)

To run ssCENTIPEDE. we performed CENTIPEDE predictions using the total number of DNA cuts in the vicinity of each motif instance as input instead of the positions of the DNA cuts. This strategy informs motif accessibility while omitting positional patterns that can be used by CENTIPEDE as a signature of TF binding. We ran CENTIPEDE similarly as described above, with the only difference being that we summed across the rows of the input Tn5 cuts matrix to generate a one-column matrix containing the total number of Tn5 cuts in the motif vicinity. This ensures that the positional information (*i.e.* where the transposition events occur relative to the motif) is omitted from CENTIPEDE.

DNase2TF

In order to run DNase2TF (v. 1.0) on ATAC-seq data, we offset all the cut points calculated using *paired_end_bam2split.r* by 4 bp before using them as input to the software, which was run with default parameters. We intersected the called footprints with each motif file and considered bound those motif instances that intersected a footprint with FDR <0.05.

HINT

We performed footprinting analyses with HINT-ATAC (RGT v. 1.1.1) using as input the broad ATAC-seq peaks and filtered BAM file from each sample. In their methods, the authors used MACS2 narrow peaks, but we found that they had lower performance compared to broad peaks (Fig. S4), so we used the latter for the analyses. We intersected the HINT output file with each motif file and considered bound every motif instance that intersected a footprint.

PIQ

We performed PWM scans using the *pwmmatch.exact.r* script included with PIQ (v. 1.3). BAM files were processed with *bam2rdata.r* due to an error in the code of *pairedbam2rdata.r* which prevented any of our BAM files from being processed. Footprinting was performed using the *pertf.r* script. Because PIQ performs its own PWM scans, we compared PIQ to BMO only on PWM matches that were shared between PIQ and BMO (using bedtools intersect).

Dataset downsampling

In order to compare the TF binding prediction methods across multiple sequencing depths, we uniformly downsampled BAM files using the *-s* flags of samtools view (v1.9), which downsamples files while maintaining read pairs intact (this behavior is not present in version 1.3). These downsampled files were used as input for peak calling and all other steps required prior to running each TF binding prediction method.

TF binding evaluation

We defined as true positives for a given TF all motif matches that fully intersected a ChIP-seq ENCODE conservative irreproducible discovery rate (IDR) narrow peak in the respective sample. We only analyzed TFs that had motifs in our database and at least 1,000 bound motif instances. For TFs with multiple motifs, we selected the motif with the highest total PWM information. For TFs with multiple ChIP-seq experiments, we selected the one with the highest number of bound motifs. To evaluate methods, we calculated the area under the precision-recall curve (AUC-PR), which informs the performance of the classifier in ranking bound and unbound motif instances, and the F1 score, which measures the performance of the threshold used to call bound motif instances. We did not use areas under the receiver-operator characteristic curve (AUC-ROC) given the highly skewed class imbalance between bound and unbound motifs, which makes AUC-ROCs an unreliable metric to evaluate TF binding predictions (*52*, *53*). AUC-PRs were calculated using packages ROCR (v. 1.0-7) and PRROC (v. 1.3) in R (*54*, *55*). To rank predictions, we used the -$\log_{10}$ adjusted $p$ values for BMO, the number of reported tags from HINT, the posteriors calculated by CENTIPEDE, the -$\log_{10}$ $p$ values calculated by DNase2TF, the purity score outputted by PIQ, and MACS2 -$\log_{10}$ $p$ values for motifs in peaks. F1 scores were calculated using the formula $F1 = 2 \cdot \frac{precision \cdot recall}{precision + recall}$ at the following thresholds for each method: BMO adjusted $p$ value < 0.05, CENTIPEDE posterior ≥ 0.99, any motif instance overlapping a HINT predicted footprint, any motif instance overlapping a DNase2TF predicted footprint with FDR value < 0.05, and any motif instance called bound by PIQ.

Mixture models for f-VICE distributions

High and low f-VICE distributions were calculated using the R package mixtools (v. 1.1.0) (*56*) using as input the normalized f-VICEs for each ATAC-seq sample, after removing low signal motifs, where the number of predicted bound instances for the motif was contained in the lowest decile of that sample. We used as cutoff a posterior probability of 0.5 to split between the high and low f-VICE distributions.

FRAP/f-VICE robust regression and CTCF-Cohesin regions comparisons

To measure the correlation between FRAP recovery times and f-VICE, we performed a literature search of reported FRAP recovery times, which are referenced in Table S4. Robust linear regressions of f-VICE and FRAP recovery times were performed with the rlm function of the R package MASS (v. 7.3-50) (*57*). For each TF with FRAP recovery times, we used the f-VICE from the motif with highest total PWM information content in our database. f-VICEs for these motifs were normalized using the same linear regression model described earlier, but including all the motifs in our database (*n*=1,850). For each sample, we required that the gene corresponding to each TF had RNA-seq TPM ≥ 1 in a related tissue in GTEx (except for pancreatic islets, where we used the RNA-seq data from (*21*)).

CTCF-Cohesin regions in GM12878 were obtained by selecting CTCF motifs that intersected conservative IDR GM12878 CTCF ChIP-seq peaks (ENCODE accessions

ENCFF096AKZ, ENCFF710VEH, and ENCFF963PJY) and the merged GM1287 RAD21 optimal IDR peaks (ENCODE accessions ENCFF753RGL and ENCFF002CPK). CTCF regions without cohesin were obtained similarly as above, but removing CTCF motifs that intersected any of the GM12878 RAD21 ChIP-seq peaks. All operations were performed with bedtools (v. 2.26.0). The choice of optimal IDR peaks for RAD21 aimed to increase the number of RAD21 peaks are included in the CTCF-cohesin$^+$ regions, therefore increasing stringency of the comparisons. We performed a quantile-based downsampling approach to make the CTCF/cohesin$^+$ and CTCF/cohesin$^-$ regions comparable regarding ChIP-seq signal, ATAC-seq signal, and FIMO motif scores. This was done by selecting all CTCF motifs encompassing the CTCF/cohesin$^+$ and CTCF/cohesin$^-$ regions and, for each feature (ATAC-seq fragments, ChIP-seq signal, or motif scores), calculating quantiles (*n*=20). Then, for every quantile, we counted the number of motifs belonging to the CTCF/cohesin$^+$ and CTCF/cohesin$^-$ regions and randomly downsampled the group with more motifs instances to have the same number of motifs as the other in that quantile. This ensured that both regions had the same number of motifs and comparable distributions of ATAC and ChIP signals and motif scores (as an example of this normalization, refer to Fig. S13A).
Pseudocode:

```
for feature in {ATAC, ChIP, PWM}:
  split feature in 20 quantiles
    for quantile in {1..20}:
      set1 = CTCF/cohesin+ ∈ feature_quantile
      set2 = CTCF/cohesin- ∈ feature_quantile
      smallest_set = smallest(set1, set2)
      largest_set = largest(set1, set2)
      n = size(smallest_set)
      randomly select n items from largest_set
```

For the main figures, we used CTCF and RAD21 experiments ENCFF963PJY and ENCFF753CPK, respectively (the same comparisons using the other CTCF/RAD21 datasets are presented in Fig. S14). The quantity labeled as relative chromatin information enrichment in Fig. 2B corresponds to the sum of positive chromatin information enrichment (above dashed line) in each V-plot, divided by the CTCF/cohesin$^-$ value for normalization.

Clustering

Chromatin information enrichment Z-score clusters were obtained using the R *k*-means implementation, using parameters *k*=3 and 1,000 random starts (Fig. S15E). Cross-tissue clustering and dendrograms were calculated using the Euclidean distances of the pairwise Spearman correlation of f-VICEs across samples. Normalized f-VICE values were converted to motif-wise Z scores before clustering.

MNase-seq data processing

Paired-end MNase unmapped reads from the lymphoblastoid cell line GM19238 were obtained from SRA, under accession SRR452483 (*58*). Reads were mapped to the hg19 reference using BWA and processed in an identical fashion to the ATAC-seq data, with an additional step to retain only sequenced fragments of length 147±2 bp, therefore enriching for mononucleosomal fragments. The MNase aggregate signal plots were generated using ngsplot (github.com/shenlab-sinai/ngsplot). For each motif plot, we used for input the BED files corresponding to the regions that were used to generate the corresponding V-plot. Motif MNase Z-scores for the clustering analyses were calculated using the MNase reads per million mapped reads (RPM) signal tracks outputted by ngsplot and the formula $Z(x) = \frac{x - mean(x)}{SD(x)}$. MNase/CIE correlations were calculated using positions ≤150 bp from the motif center.

Chromatin information enrichment asymmetry

Chromatin information enrichment asymmetry was calculated as the $\log_2$ ratio between the positive information content enrichment in the left and right of the motif center. To estimate significance, we used a permutation test where each fragment midpoint had a 50% chance of changing its direction relative to the motif while keeping the same distance (*i.e.* multiply its x-axis value by -1). We calculated the asymmetry of the permuted V-plots ($n$ = 100,000) to generate a null distribution of asymmetry. Because the null was normally distributed based on Kolmogorov-Smirnof and Shapiro normality tests, we were able to estimate $p$ values beyond the number of permutations by calculating the observed asymmetry Z-score relative to the null distribution. To calculate the nearest TSS directionality bias, we counted the number of active protein-coding TSS (GENCODE V19) (determined with the presence of LCL Cap analysis gene expression (CAGE) tag clusters, described in the next session) on either side of the motif and calculated the $\log_2$ ratio of the two. For the proximal and distal motif V-plots, we restricted our analyses to motifs occurring ≤1kb or ≥10kb from the nearest CAGE-supported TSS of any type (*e.g.* lincRNAs, pseudogenes; GENCODE V19). Enrichments of the plots in Fig. 2F were calculated by randomly permuting the signal of the points in the x- and y-axis ($n$=10,000 permutations).

CAGE tag cluster identification

We downloaded CAGE data (fastq files) for 154 LCL samples (*59*) and mapped to hg19 using STAR (version 2.5.4b; default parameters) (*60*) and pruned the mapped reads to high quality reads (using samtools view v. 1.3.1; options *-F 4 -q 255*). We used the paralu method (*61*) to identify clusters of CAGE start sites (CAGE tag clusters). We called TCs in each individual sample using raw tag counts, requiring at least 2 tags at each included start site and allowing single base-pair tag clusters ('singletons') if supported by >2 tags. We then merged the tag clusters on each strand across samples. For each resulting segment, we calculated the number of LCL samples in which TCs overlapped the segment. We included the segment in the consensus TCs set if it was supported by independent TCs in at least 10 individual LCL samples, resulting in n=10 tag clusters. We then filtered out regions blacklisted by the ENCODE consortium due to poor mappability using bedtools (v. 2.26.0) to obtain the final set of LCL tag cluster regions.

DNA binding domain enrichments

DNA binding domains (DBD) enrichments were performed using a f-VICE rank sum permutation test. We assigned DBDs to the non-redundant motifs that mapped between our database and the one reported in (*62*), which has manually curated DBD-motif assignments. In order to map motifs between databases, we used tomtom (*63*) and selected motif matches with $p$-value < 0.05 after a conservative Bonferroni adjustment using all comparisons as denominator (*i.e.* number of motifs in our database times the number of motifs in the queried database), which yielded high-confidence DBD assignments for 402 of 540 motifs. We used the f-VICE rank from each motif to calculate the f-VICE rank sum the DBD and compared the observed value to a null distribution of 100,000 rank sums obtained from randomly permuting gene labels. This approach ensures that all the DBD retain their sizes during each permutation. We retained DBDs with at least 5 motifs and calculated the f-VICE enrichments for each DBD using the $\log_2$ of observed f-VICE rank sum divided by the median of the null. FDR was calculated separately per sample, using the empirical *p*-value from the 100,000 permutations. We simultaneously performed a similar analysis using InterPro protein domains (v. 72) (*64*) (Fig. S18). In order to assign domains to motifs, we first mapped our motifs to CIS-BP database (Build 1.02) (*65*), which has high-confidence motif-gene assignments, and retained genes that mapped to a single motif using the same approach described above. Each gene was then linked to a motif f-VICE score (n = 475) and we only retained domains with at least 5 genes after motif-gene mapping. Permutation and enrichments were calculated identically as described above.

cis-eQTL enrichments

Feature enrichments in eQTLs were calculated using GREGOR (*66*) and QTL tools fenrich (*67*). We used the lymphoblastoid cell line (LCL) eQTLs sets from Geuvadis (*32*) and GTEX (*31*) (FDR<5%). GREGOR background estimations were performed using SNPs with LD 0.99 for eQTL, with a maximum distance of 1 Mb from the variants of interest. Variants used as input for GREGOR were pruned to have maximum linkage disequilibrium $r^2$ of 0.8 with any other variant. For fenrich, we used the most significant SNP per gene as input.

ATAC-seq allelic imbalance analyses

To determine SNP allelic bias in ATAC-seq data, we used the publicly available data from Buenrostro *et al*, listed in Table S1, the Parker lab GM12878 sample discussed here, and the ABCU196 islet sample introduced in (*21*). For GM12878 data, adapters were trimmed using cta (v. 0.1.2), and reads mapped to hg19 using bwa mem (default options except for the -M flag). Bam files were filtered to high-quality autosomal read pairs using samtools view (-f 3 -F 4 -F 8 -F 256 -F 2048 -q 30; v. 1.3.1). WASP (v. 0.2.1, commit 5a52185; python version 2.7) (*68*) was used to diminish reference bias; for remapping the reads as part of the WASP pipeline, we used the same mapping and filtering parameters described above for the initial mapping and filtering. Duplicates were removed using WASP's rmdup_pe.py script. We used the phased GM12878 VCF file downloaded from ftp://ftp-trace.ncbi.nlm.nih.gov/giab/ftp/release/NA12878_HG001/NISTv3.3.1/GRCh37/HG001_GRCh37_GIAB_highconf_CG-IllFB-IllGATKHC-Ion-10X-SOLID_CHROM1-X_v.3.3.1_highconf_phased.vcf.gz. To avoid double-counting alleles, overlapping read pairs were clipped using bamUtil clipOverlap (v. 1.0.14; genome.sph.umich.edu/wiki/BamUtil:_clipOverlap). For the Buenrostro *et al* data, the bam files from the samples in Table S1 were then merged to create a single GM12878 bam file using samtools merge (v. 1.3.1). For each heterozygous autosomal SNP, we then counted the number of reads containing each allele, using only bases with base quality of at least 20. We used a two-tailed binomial test that accounted for reference allele bias to evaluate the significance of the allelic bias at each SNP (as described in (*21*); when calculating the expected fracRef, SNPs in the top 25th percentile of read coverage were downsampled to the 50th percentile coverage and SNPs with coverage less than 10 were excluded). When performing the binomial test, we downsampled the coverage at each SNP such that each SNP had coverage = 20 (to reduce coverage-related biases). The islet ATAC-seq data was processed and tested as described in (*21*), except that we also downsampled coverage at each SNP to 20 reads when performing the binomial test. We did not test SNPs in regions blacklisted by the ENCODE Consortium because of poor mappability (wgEncodeDacMapabilityConsensusExcludable.bed and wgEncodeDukeMapabilityRegionsExcludable.bed). We retained for downstream analyses all loci with nominally significant binomial $p$ values ($p < 0.05$) and at least 2 reads (10%) mapped to any allele. After selecting loci that intersected at least one predicted bound 6-mer (see next section), we obtained 257, 1,449, and 15,802 imbalanced SNPs for the two GM12878 datasets (this study and Buenrostro) and pancreatic islets, respectively.

6-mer f-VICE calculations

We generated a list of all possible DNA 6-mers and scanned the hg19 reference genome to obtain the coordinates for all their corresponding matches. Similarly to motifs, we only retained 6-mer matches that were in mappable regions and did not intersect blacklisted regions. For each set of 6-mer matches, we used BMO to determine the subset that was predicted bound. We calculated the normalized f-VICE for each 6-mer using exactly the same steps as in the motifs, including using linear regression to control for chromatin accessibility. For each 6-mer, we determined its immediate neighbors in sequence space (every 6-mer that differed by exactly 1 letter; Hamming distance = 1) and calculated the f-VICE differences between each of the neighbors relative to the original 6-mer. We then calculated the Euclidean distance between the neighbors with highest and lowest f-VICE to determine what was the f-VICE range associated with that 6-mer.

For each locus with significant allelic imbalance, we calculated the f-VICE associated with all six 6-mer instances overlapping with each allele. For each sample, we determined the f-VICE decile changes associated with every SNP tested for allelic imbalance and determined the matrix of the $\log_2$ ratios of each possible decile change in the imbalanced versus all tested SNPs. To test for significance, we devised a permutation test where all symmetrical pairs of f-VICE deciles, $x_{i,j}$ and $x_{j,i}$ for $i, j \in \{1,2, \dots ,10\}$, had a 50% chance of switching their $\log_2$-ratio values in each permutation ($n$ = 1,000,000 permutations).

Data availability

Code and scripts used for the analyses performed in this study are publicly available at http://github.com/ParkerLab/chromatin_information. BMO and atactk are publically available at http://github.com/ParkerLab/BMO and http://github.com/ParkerLab/atactk.

**Supplementary text**

Transcription factor binding prediction methods comparison

BMO builds on previous reports that the degree of chromatin accessibility around a motif (*16–18*) and the presence of co-occurring motifs (*19*) positively correlates with TF binding, and uses TF-specific negative binomial models of these two signals to estimate the likelihood of a bound instance (*1*). We benchmarked the performance of BMO and other unsupervised TF binding prediction algorithms using ATAC-seq datasets from the GM12878 and HepG2 (*48*) cell lines and their corresponding TF ChIP-seq data ($n$=41 and $n$=59, respectively; Table S1). We compared BMO to three footprinting-based algorithms (HINT-ATAC (*14*), DNase2TF (*13*), PIQ (*12*)), to CENTIPEDE, which learns informative DNA cut patterns indicating TF binding (*20*), and to a baseline classifier that labels TF motifs within ATAC-seq peaks as bound. To evaluate methods, we calculated the area under the precision-recall curve (AUC-PR), which informs the performance of the classifier in ranking bound and unbound motif instances, and the F1 score, which measures the performance of the threshold used to call bound motif instances.

BMO outperformed all methods in our high-signal GM12878 dataset (Fig. 1E), whereas BMO and CENTIPEDE had similarly high performance in lower-signal datasets (Figs. S1, S3). DNase2TF had lower performance in the lower-signal datasets (Fig. S3). PIQ cannot use custom TF motif scans and therefore required separate benchmarking, which revealed lower performance compared to BMO (Fig. S5). These results were consistent across ATAC-seq replicates and cell lines, including downsampled data representing shallower sequencing depths (Fig. S6). Of note, the AUC-PR of footprinting-based methods was lower overall due to their inability to classify motifs occurring outside ATAC-seq peaks, which we reason contain true TF binding sites and negatively affect PR-AUCs. While their F1 scores indicate that this effect is less pronounced when taking into account the thresholds to call bound motif instances, their performance was still consistently lower than non footprinting-based methods (Fig. S3). Overall, the two footprinting-agnostic methods (BMO and CENTIPEDE) outperformed footprint-based methods on the majority (median of 81% across datasets) of tested TFs. These results indicate that TF binding is more accurately predicted using a simple chromatin accessibility model tuned to each TF motif.

We next sought to determine if the CENTIPEDE approach relied on spatial DNA cut patterns, or if the overall accessibility in the region was sufficient for high performance. We devised an alternative implementation of CENTIPEDE that ignores the DNA cut positions (signal-sum CENTIPEDE; ssCENTIPEDE) and masks any footprint-like patterns, but not the chromatin accessibility in the region (1). This ssCENTIPEDE approach performed almost identically to CENTIPEDE (Figs. S7, 1E), again indicating that footprint patterns in chromatin profiles are not necessary for high prediction performance. One corollary expectation from this conclusion is that footprint-based algorithms should perform comparatively worse when predicting binding for TFs with a low impact on local chromatin. To test this, we compared performance across f-VICE tertiles representing low (tertile one), intermediate (tertile two), and high (tertile three) f-VICEs. Notably, BMO and (ss)CENTIPEDE had relatively higher performance on lower f-VICE tertiles one and two (Figs. 1E, S3). Our findings indicate that footprinting-based methods are more sensitive to the local TF-chromatin architecture.

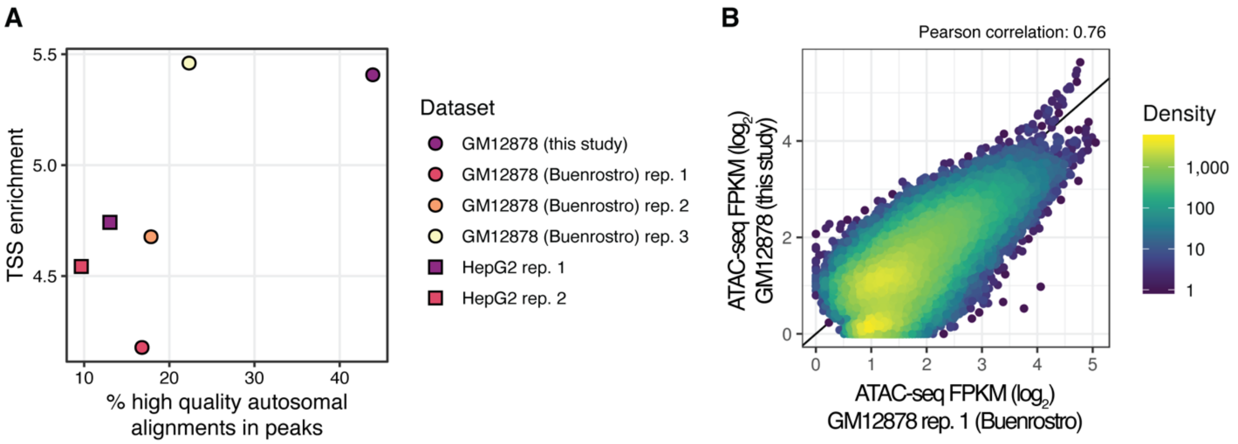


**Fig. S1. ATAC-seq datasets signal-to-noise comparisons.**

(**A**) Scatter plots of the percent high quality autosomal alignments (%HQAA) in ATAC-seq peaks distribution and TSS enrichments of GM12878 and HepG2 datasets, obtained using Ataqv (github.com/ParkerLab/ataqv). (**B**) Scatter plot of the ATAC-seq signal in the union of the MACS2 broad peaks called in the two GM12878 datasets. Each point corresponds to one ATAC-seq peak. Solid diagonal line, identity ($x$=$y$).

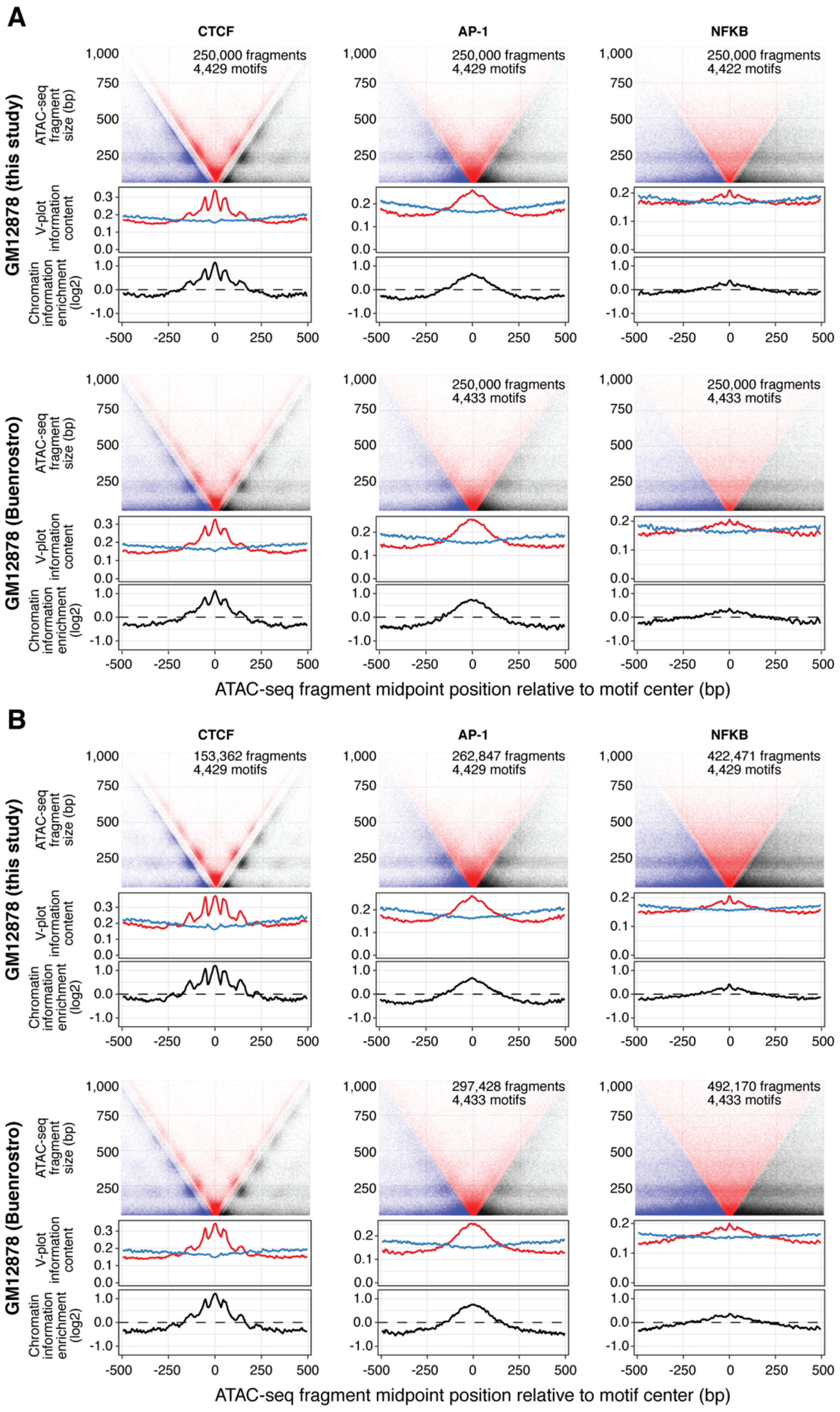
A
CTCF
AP-1
NFKB
GM12878 (this study)
GM12878 (Buenrostro)
250,000 fragments
4,429 motifs
4,422 motifs
4,433 motifs
ATAC-seq fragment size (bp)
V-plot information content
Chromatin information enrichment (log2)
ATAC-seq fragment midpoint position relative to motif center (bp)
B
153,362 fragments
262,847 fragments
422,471 fragments
297,428 fragments
492,170 fragments

**Fig. S2. GM12878 V-plots.**
(**A**) V-plots for the same TFs as in Figure 1B across GM12878 datasets. Upper: ATAC-seq fragment distribution. Middle: observed (red) and expected (blue) information content tracks, used to calculate chromatin information enrichments (bottom). V-plots were downsampled to equal number of ATAC-seq fragments and motifs between TFs by selecting the top *n* motifs, ranked by number of ATAC-seq fragments, and then further downsampling to 250,000 fragments. *n* represents the smallest number of bound motifs among the plotted TFs per sample. (**B**) Similar to (A), but randomly downsampling to exactly *n* motifs (without ranking by signal or further downsampling the number of ATAC-seq fragments). This was performed to demonstrate that the differences in chromatin architecture are intrinsic to the TF and evident regardless of downsampling method.

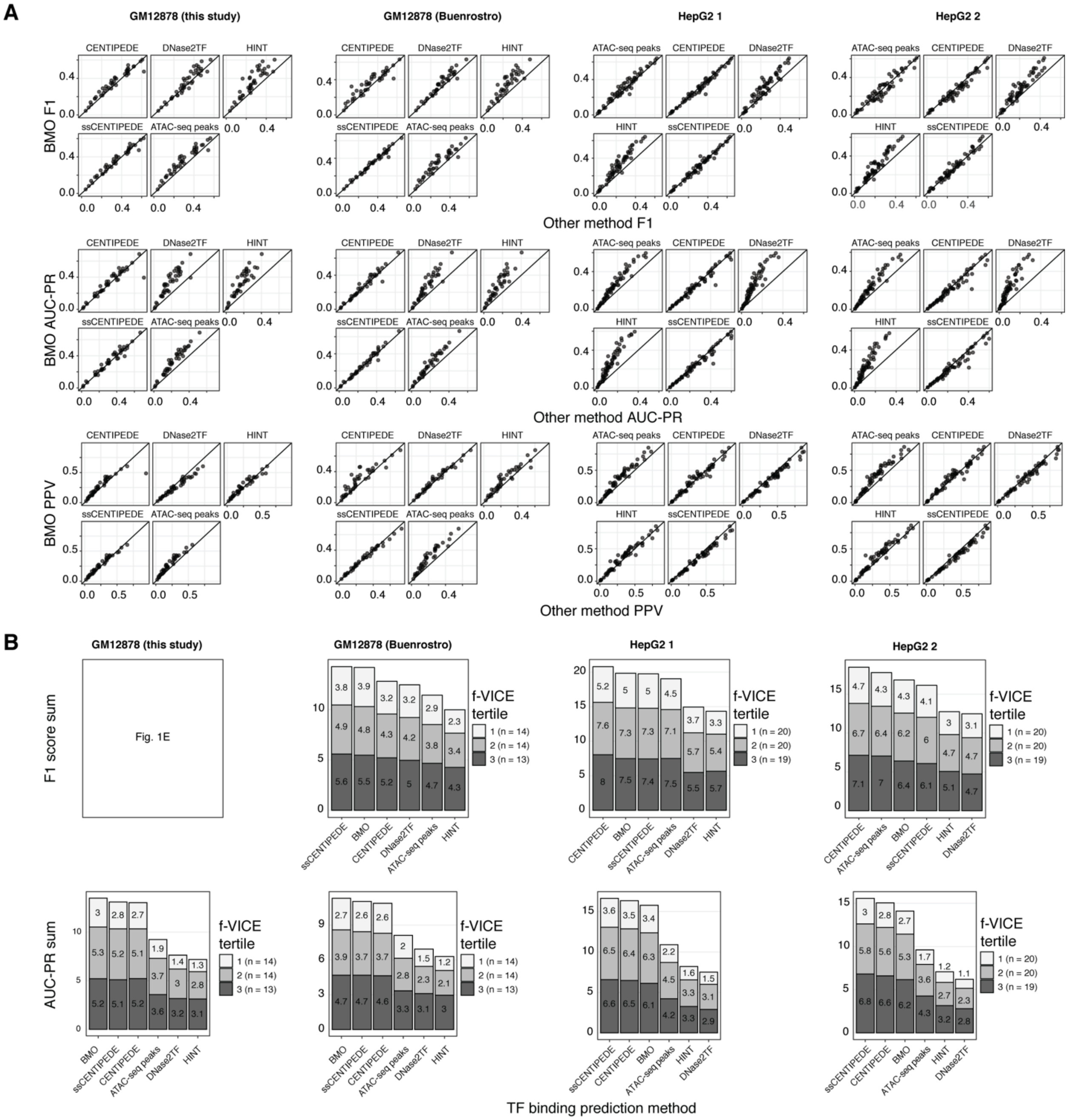


**Fig. S3. TF binding prediction methods comparisons across datasets.**

(**A**) F1, positive predictive value (PPV), and AUC-PR scatter plots of BMO versus other TF binding prediction methods across multiple ATAC-seq datasets. Each point corresponds to a TF with ChIP-seq data. (**B**) Total F1-score and AUC-PR across datasets, separated into f-VICE tertiles. Solid diagonal line, identity (*x*=*y*).

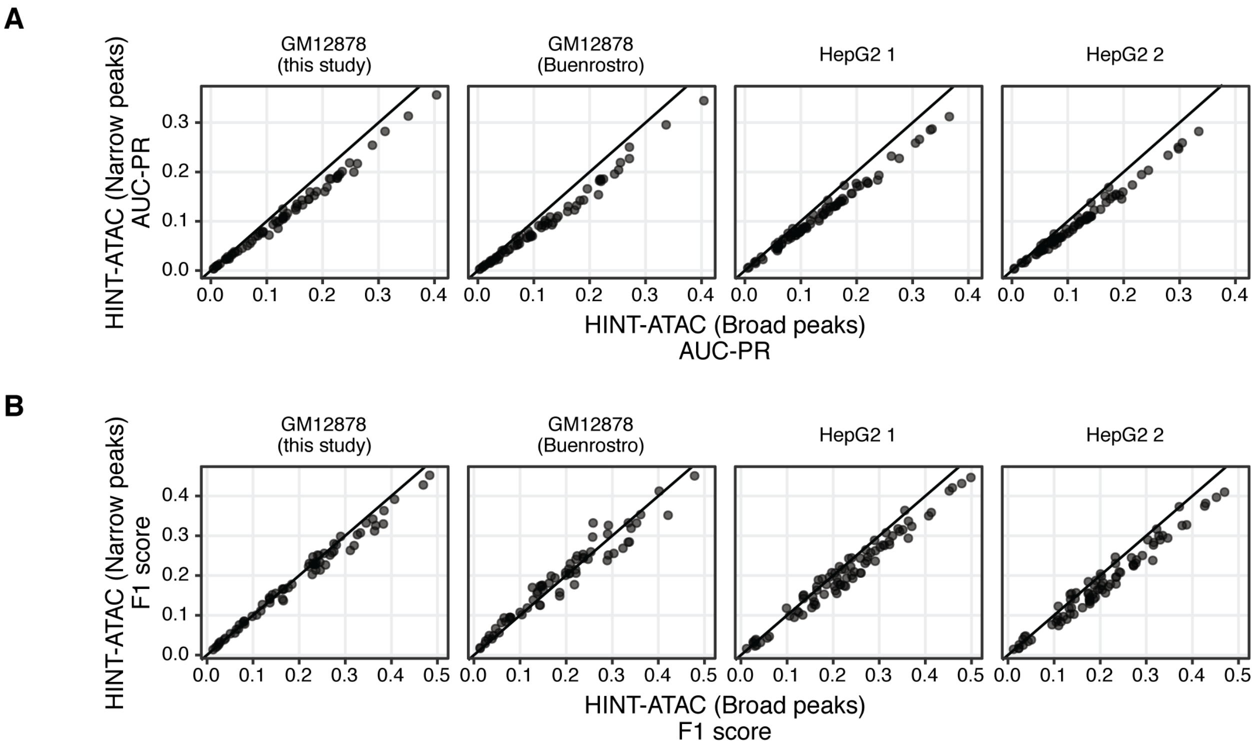


**Fig. S4. HINT-ATAC performance using narrow or broad peak calls.**

Scatter plots of AUC-PRs (**A**) and F1 scores (**B**) across datasets. Each point corresponds to a TF with ChIP-seq data. Solid diagonal line, identity (*x*=*y*).

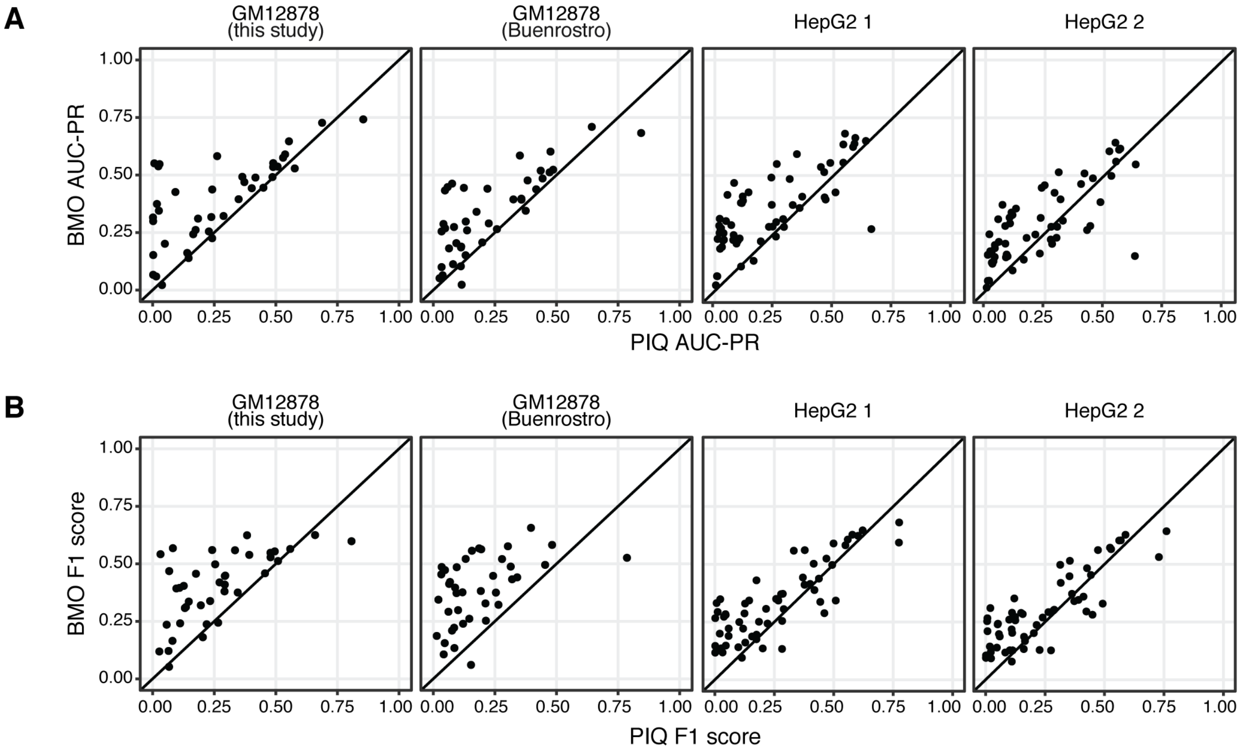


**Fig. S5**. **BMO and PIQ comparisons.**
Scatter plots of AUC-PR (**A**) and F1 scores (**B**) across datasets comparing BMO and PIQ. Each point corresponds to a TF with ChIP-seq data. Solid diagonal line, identity (*x*=*y*).

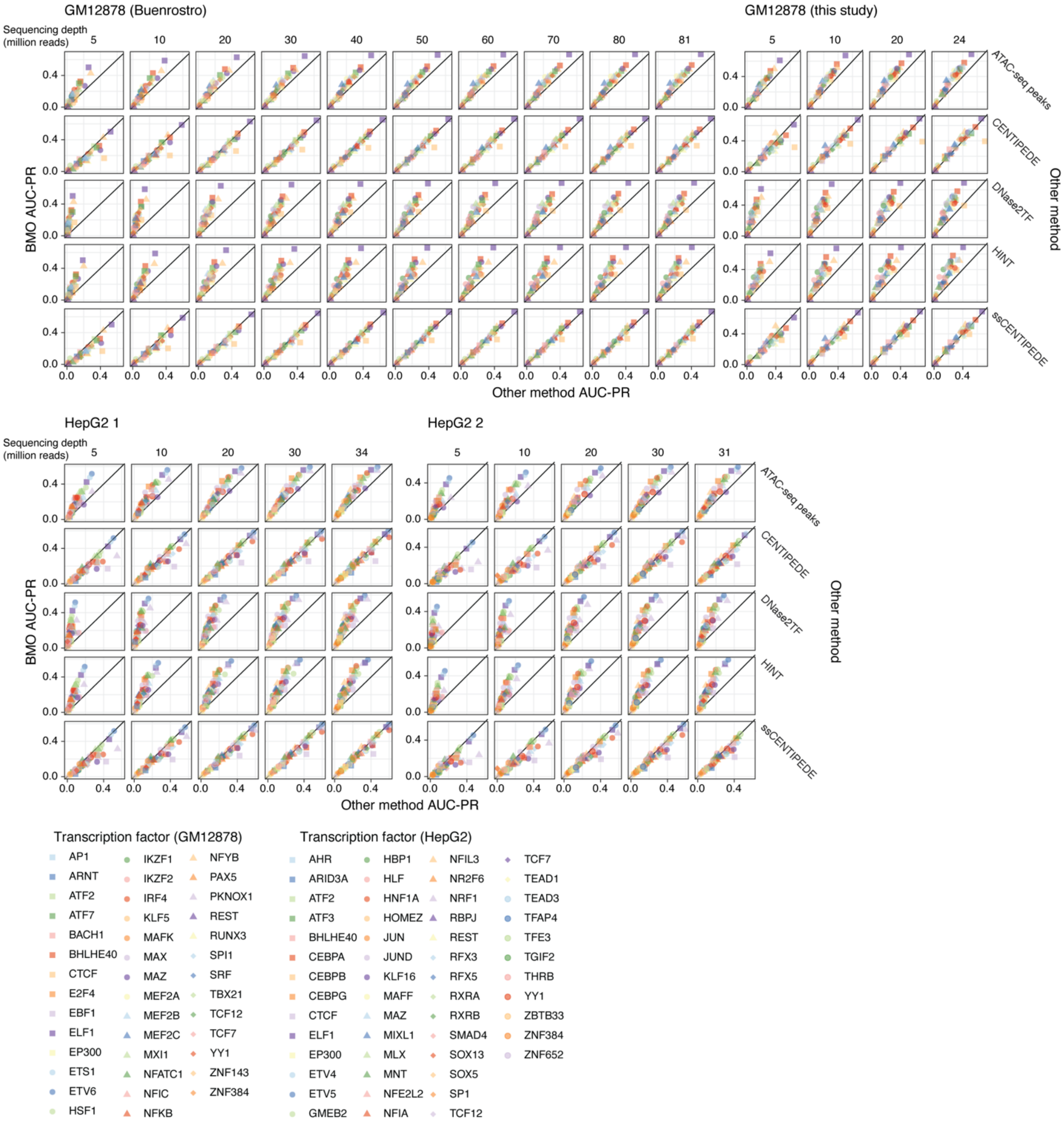


**Fig. S6**. **TF binding prediction methods comparisons across sequencing depths.** AUC-PR scatter plots of BMO versus other methods across different sequencing depths, shown in millions of reads in the top of each facet column. Each point corresponds to one TF with ChIP-seq data. Solid diagonal lines, identity (*x*=*y*).

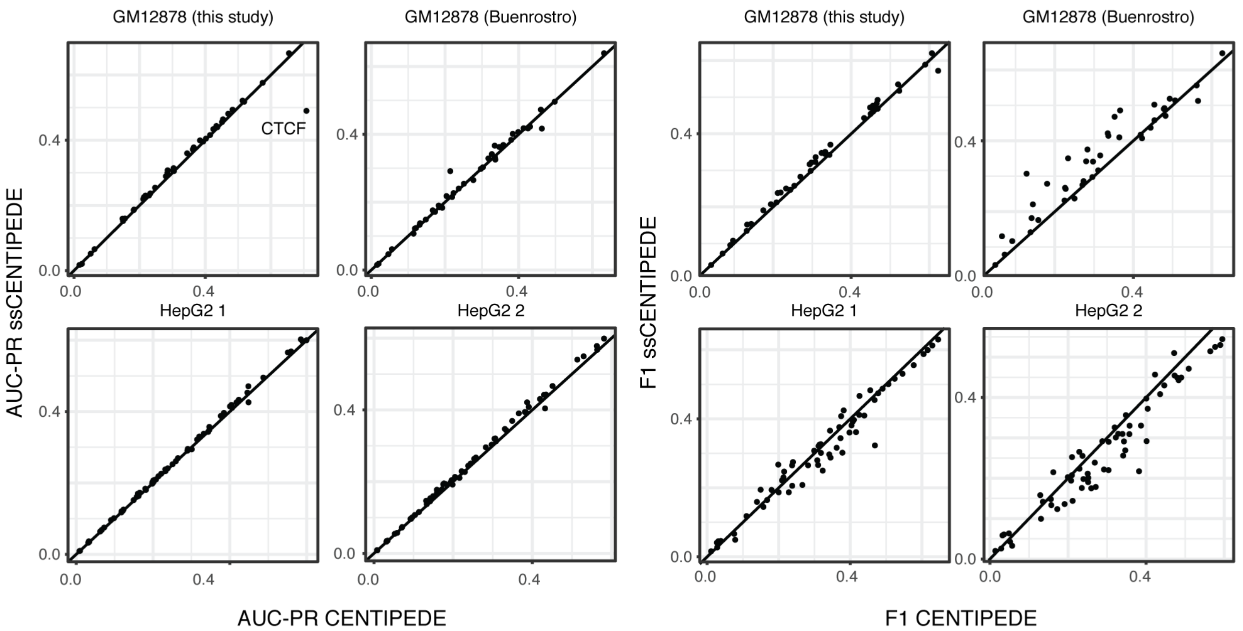


**Fig. S7. CENTIPEDE and ssCENTIPEDE perform similarly across datasets.**
Scatter plots of ssCENTIPEDE and CENTIPEDE AUC-PRs and F1 scores across multiple datasets. Each point corresponds to one TF with ChIP-seq data. Solid diagonal line, identity (*x*=*y*).

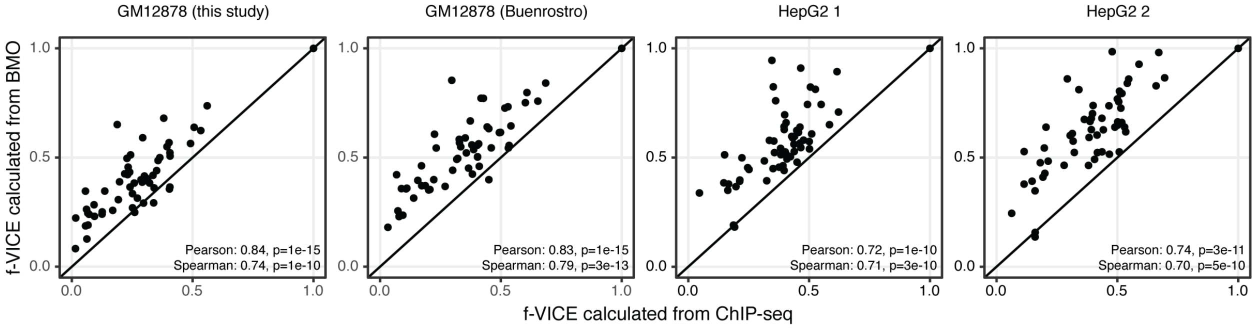


**Fig. S8**. **BMO and ChIP-seq f-VICEs are correlated.**

Correlation of f-VICEs calculated from BMO predictions and from the respective ChIP-seq data across ATAC-seq datasets. Note that BMO f-VICEs are consistently higher than ChIP-seq f-VICEs. This is due to a higher number of predicted bound motif instances in BMO, which motivated us to normalize f-VICEs using the linear regression approach described in the methods (f-VICEs are not normalized using regression in this figure owing to low $n$). Solid diagonal line, identity ($x$=$y$).

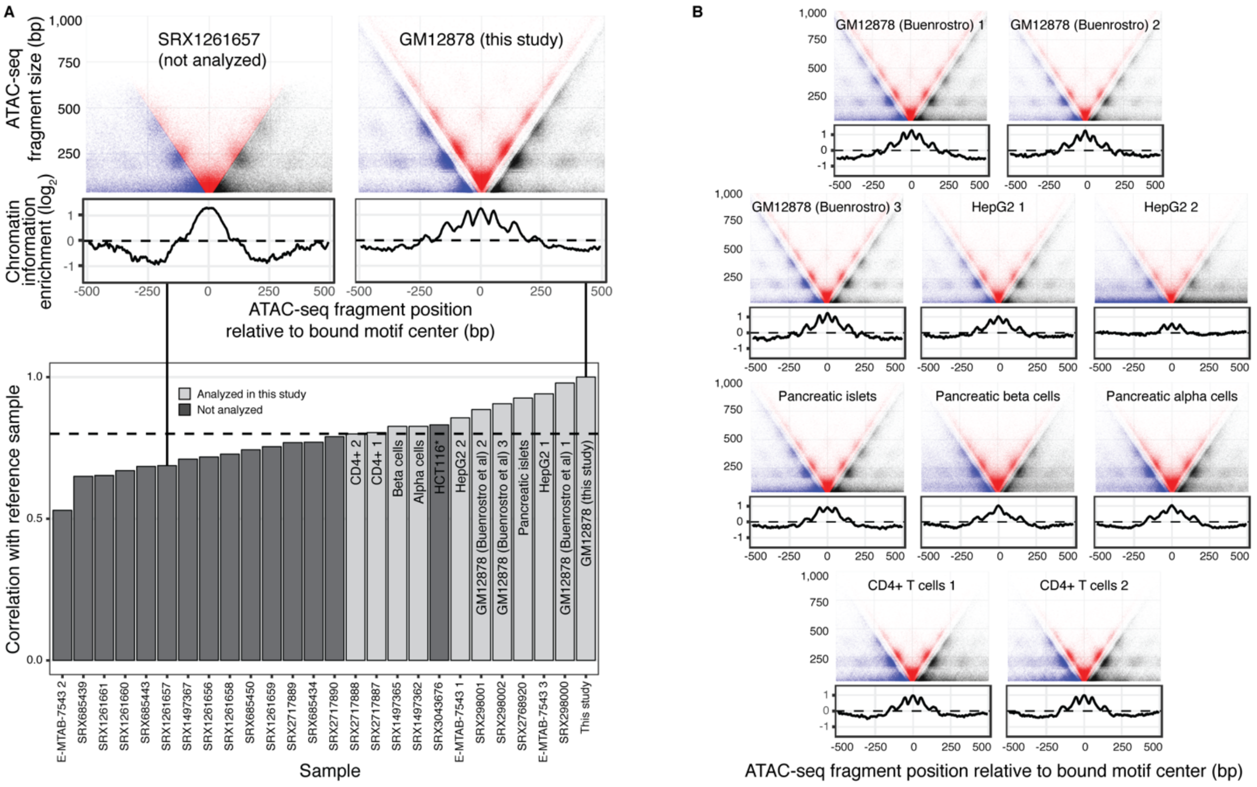


* Did not have another sample of the same tissue/cell line that passed QC

**Fig. S9. Selection of additional ATAC-seq samples using ubiquitous and conserved CTCF-cohesin binding sites.**

(**A**) Upper: examples of V-plots for the reference ubiquitous and conserved CTCF-cohesin binding sites indicating a high-quality and low-quality sample (the latter shown for exemplification purposes and not included in this study). Lower: chromatin information enrichment correlation between the CTCF-cohesin binding sites across multiple experiments to a reference sample. (**B**) V-plots of the same regions in the other samples selected for this study. Y-axes labels are the same as the upper plot in panel A.

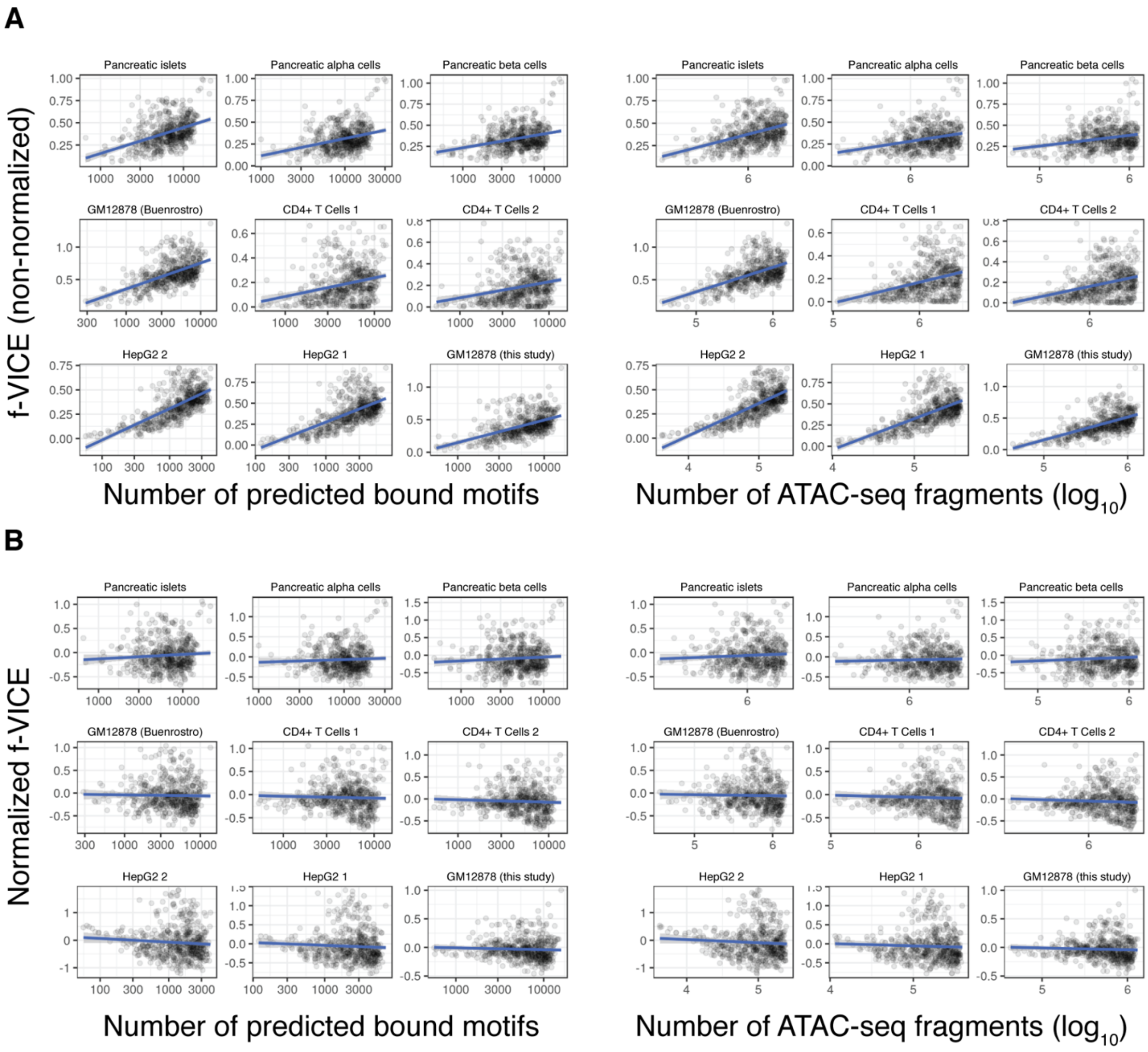


**Fig. S10. Normalization of f-VICE.**
(**A**) Scatter plots of f-VICE as a function of number of predicted bound motifs or ATAC-seq signal. (**B**) Same data after normalization using a linear regression model that accounts for both variables (described in the Methods section). Each point corresponds to a motif (*n*=540).

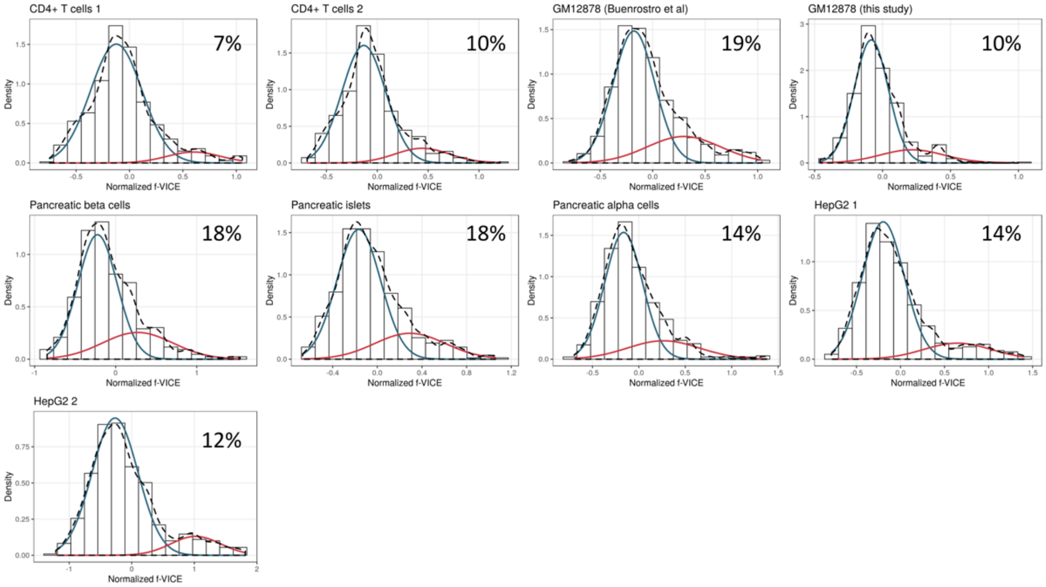


**Fig. S11. f-VICE distributions across samples.**
Histograms and density plots of the empirical (dashed) and high/low f-VICE distributions Gaussian fits (red and blue, respectively) across all the datasets surveyed in this work. Percentages in the upper right corner of plots represent the high f-VICE distribution.

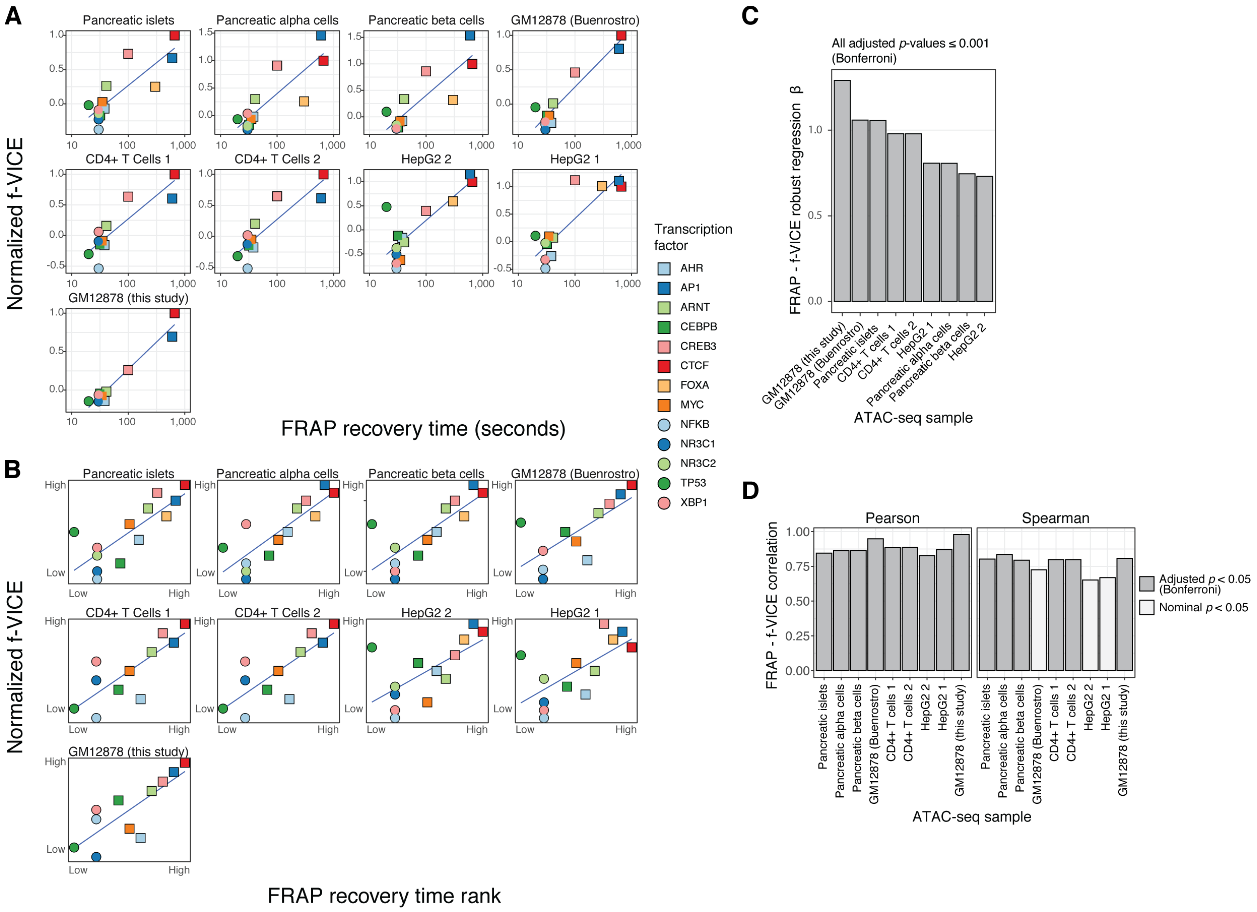


**Fig. S12. f-VICE correlation with FRAP recovery times in multiple datasets.**
(**A**) Scatter plots of mammalian FRAP recovery times and f-VICEs across multiple datasets. (**B**) Similar to (A), but showing f-VICE and FRAP ranks (similar to a Spearman correlation). Solid blue lines in (A) and (B), linear model fit. (**C**) Robust linear regression betas for the plots shown in A. D) Pearson and Spearman correlations of the plots shown in (A). All correlations were significant at $p<0.05$.

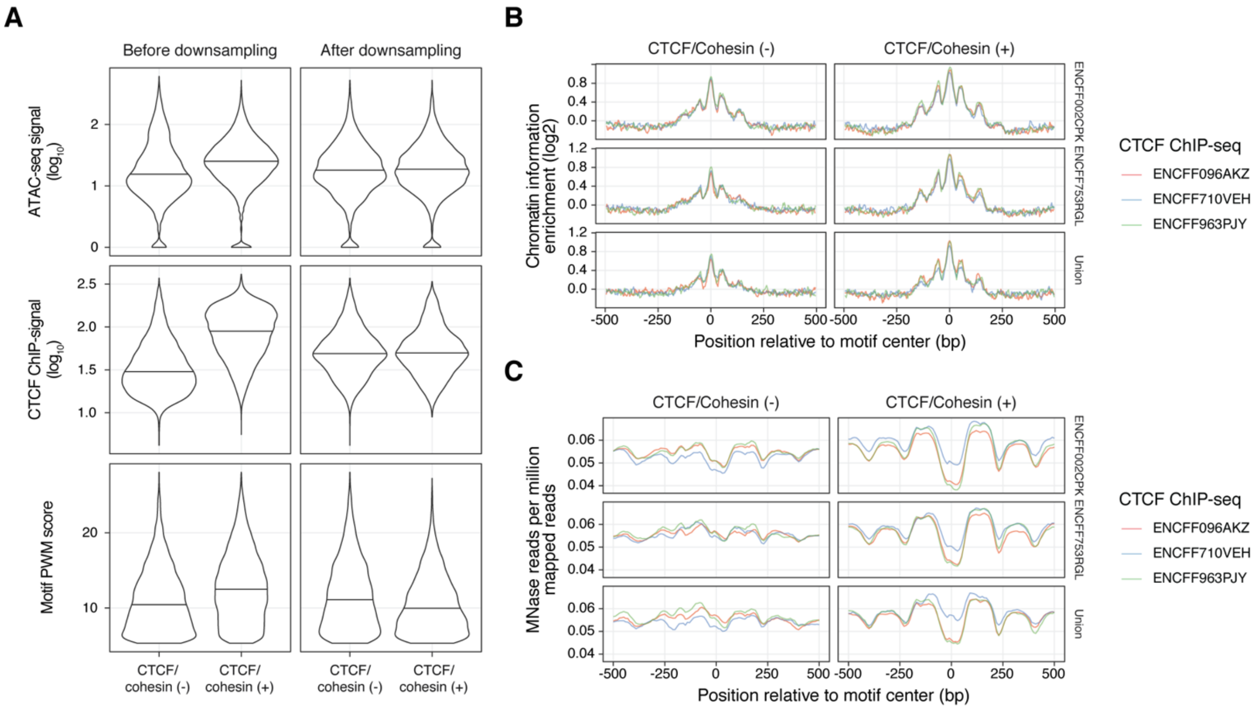


**Fig. S13. GM12878 CTCF/cohesin$^+$ and CTCF/cohesin$^-$ regions.**
(**A**) Example distributions of ATAC-seq signal, ChIP-seq signal, and motif PWM match score before and after quantile-based downsampling of the CTCF/cohesin$^+$ and CTCF/cohesin$^-$. Datasets: ENCFF963PJY (CTCF) and ENCFF002CPK (Rad21). Other CTCF/Rad21 ChIP-seq dataset combinations not shown. Horizontal lines, median. (**B**) Chromatin information tracks of CTCF/cohesin$^+$ and CTCF/cohesin$^-$ using genomic regions obtained from different GM12878 CTCF and RAD21 ChIP-seq datasets combinations. The facet labeled "Union" corresponds to the union of the two GM12878 RAD21 ChIP-seq datasets. (**C**) Corresponding MNase signal at the regions shown in (B).

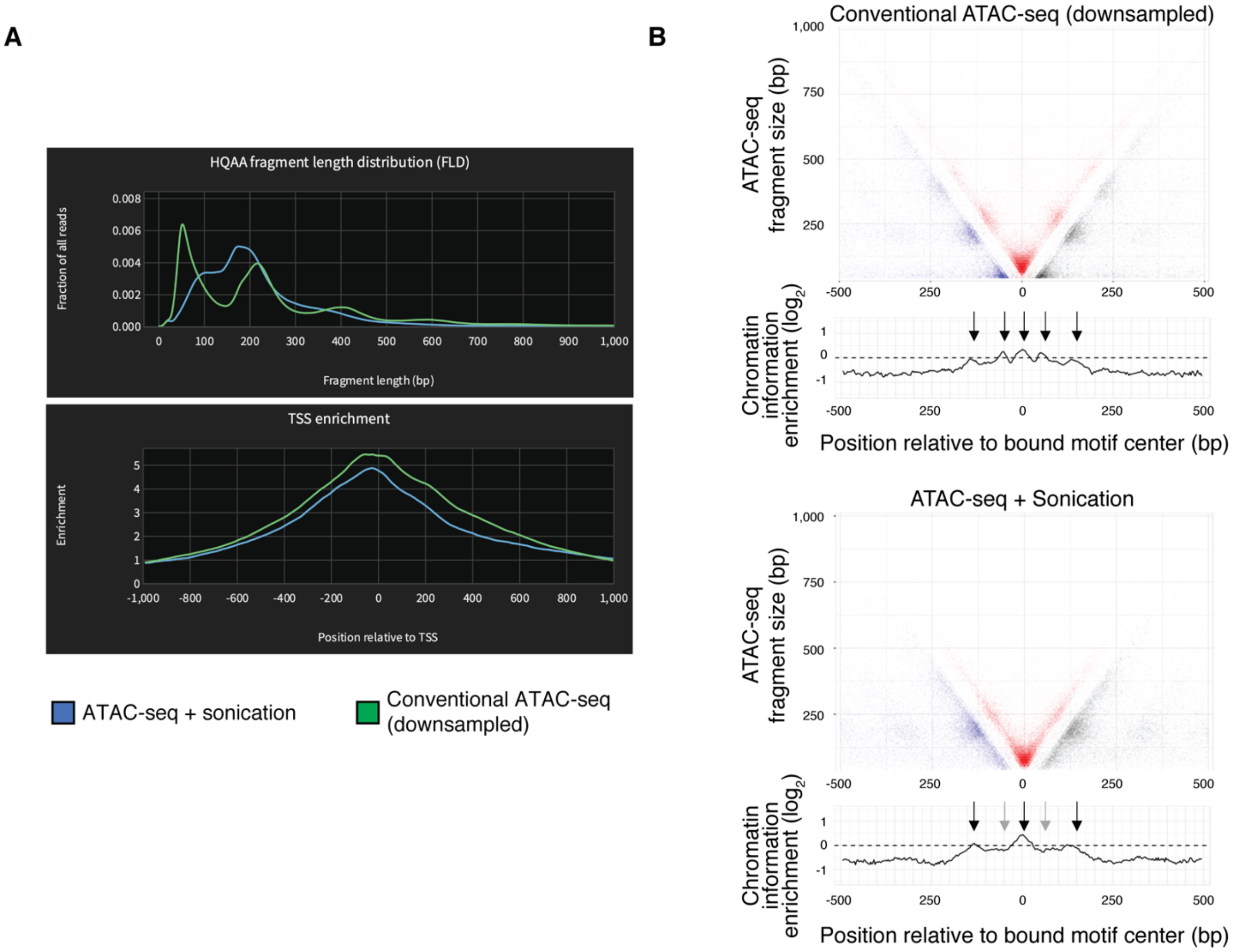


**Fig. S14**. **Sonicated GM12878 ATAC-seq data.**
(**A**) Ataqv (github.com/ParkerLab/ataqv) screenshot showing the fragment size distribution and TSS enrichments of the conventional and sonicated GM12878 ATAC-seq datasets generated in this study. HQAA = high-quality autosomal alignments. (**B**) V-plots of the reference conserved CTCF-cohesin regions in the two datasets. "Conventional ATAC-seq" refers to the sample labeled as "GM12878 (this study)" in other figures. However, this dataset was downsampled to the same depth as the sonicated dataset (3.45 million reads) for the analyses presented in this figure and Fig. 2B in order to make datasets directly comparable. Black arrows, CIE peaks in both samples. Gray arrows, CIE peaks not in the sonicated sample.

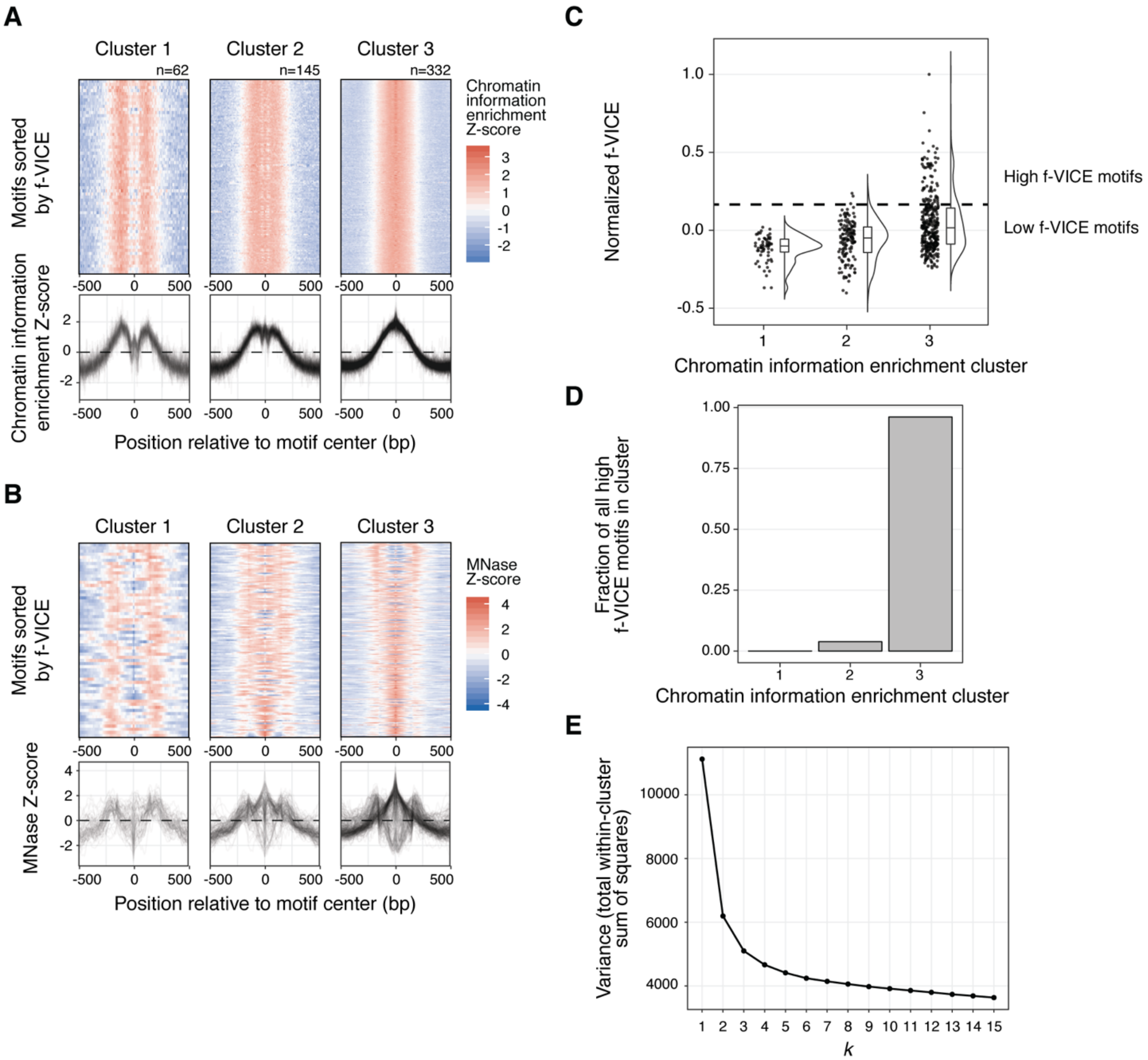


**Fig. S15**. **Chromatin information clusters in GM12878.**
(**A**) Chromatin information enrichment Z-score clusters and (**B**) their corresponding MNase Z-scores. All clusters are sorted by f-VICE on the *y*-axis. (**C**) Distribution of f-VICEs across chromatin information enrichment clusters. (**D**) Fraction of total high f-VICE TFs per chromatin information cluster (based on the mixture model distributions). (**E**) Elbow plot showing within-cluster variance for different *k* values in the chromatin information enrichment *k*-means clustering in panel (A).

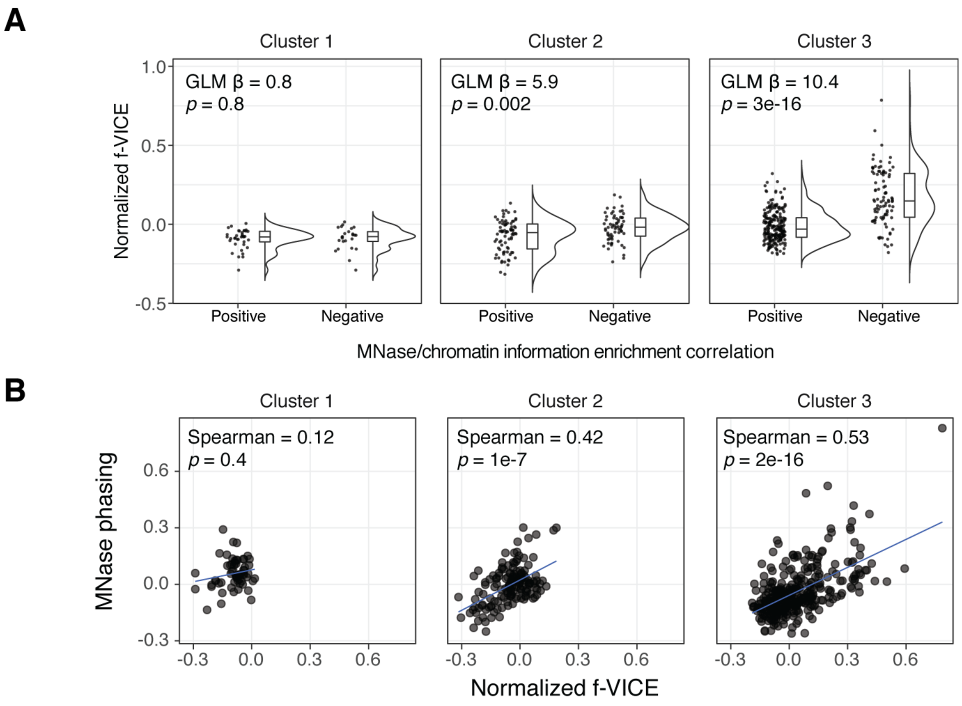


$$\text{MNase phasing} = \log_2\left(\frac{\text{MNase}_{\text{flanking}}}{\text{MNase}_{\text{motif}}}\right)$$

**Fig. S16**. **f-VICE correlation with nucleosome phasing.**
In this plot, we show two independent approaches for correlating f-VICE to nucleosome phasing. (**A**) f-VICE distributions for motifs with positive and negative correlation between chromatin information and MNase Z-scores (≤150 bp from motif center) across the different k-means clusters, labeled 1-3 in the header. GLM = generalized linear model. (**B**) Correlation between f-VICE and the $\log_2$ ratio of the MNase signal at the motif vicinity (±125-150 bp from motif center) divided by the MNase signal at the motif (±25 bp from motif center). Positives ratios indicate nucleosome phasing.

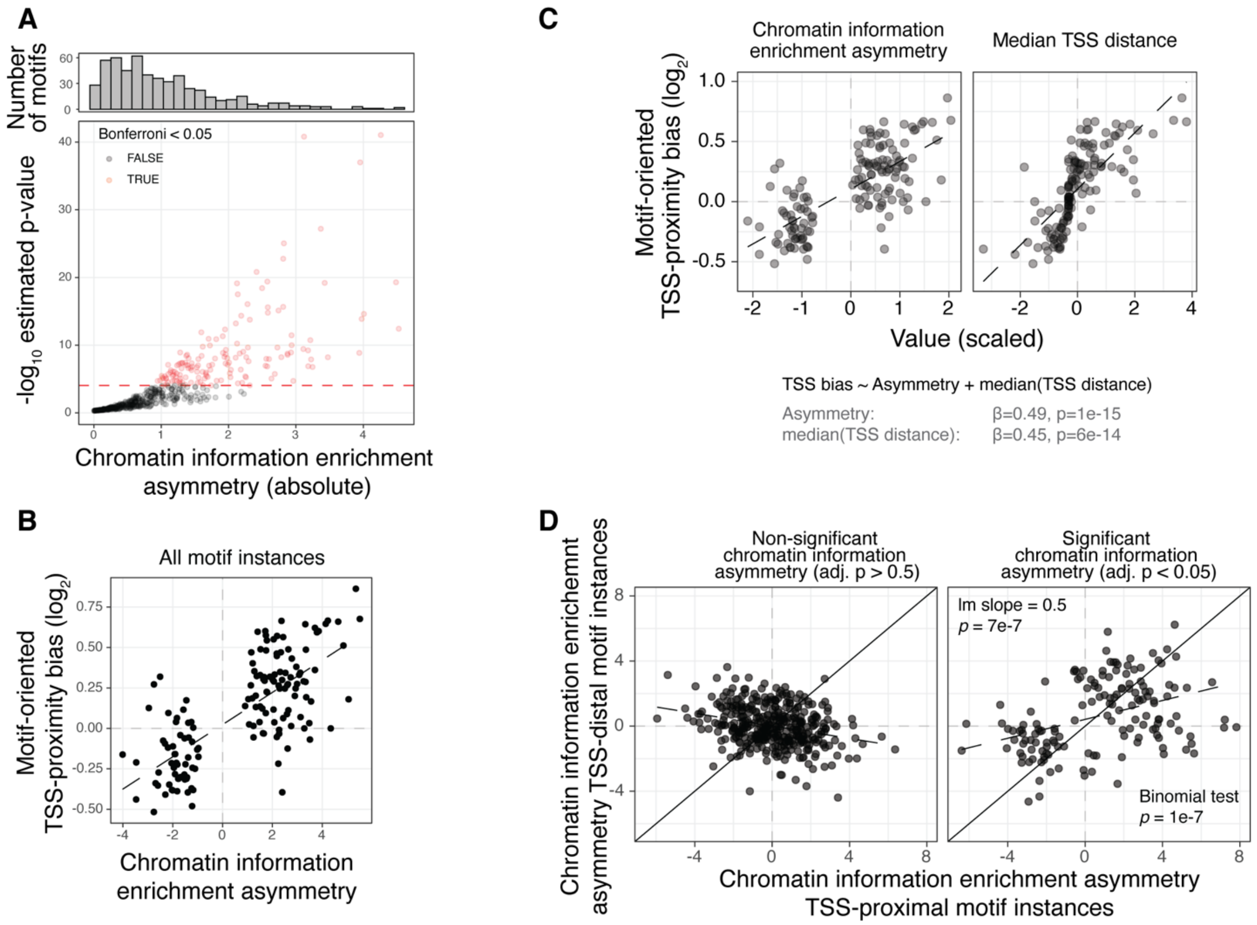


**Fig. S17**. **Motifs with information asymmetry in GM12878.**
(**A**) Chromatin information asymmetry distribution in GM12878. Red dashed line represent the Bonferroni *p*-value cutoff threshold. (**B**) Relationship between chromatin information asymmetry and motif-oriented TSS proximity bias based on all motif instances. (**C**) Motif-oriented TSS proximity bias chromatin as a function of information asymmetry and median nearest TSS distance. We performed a regression analysis of nearest TSS direction bias and chromatin information enrichment asymmetry, controlling for TSS distance (*1*). Chromatin information enrichment asymmetry remained significant when controlling for TSS distance. (**D**) Concordance of chromatin information asymmetry direction between TSS-distal and TSS-proximal motif instances. Solid diagonal line, identity ($x$=$y$). Dashed black lines, linear model (lm) fit in the data.

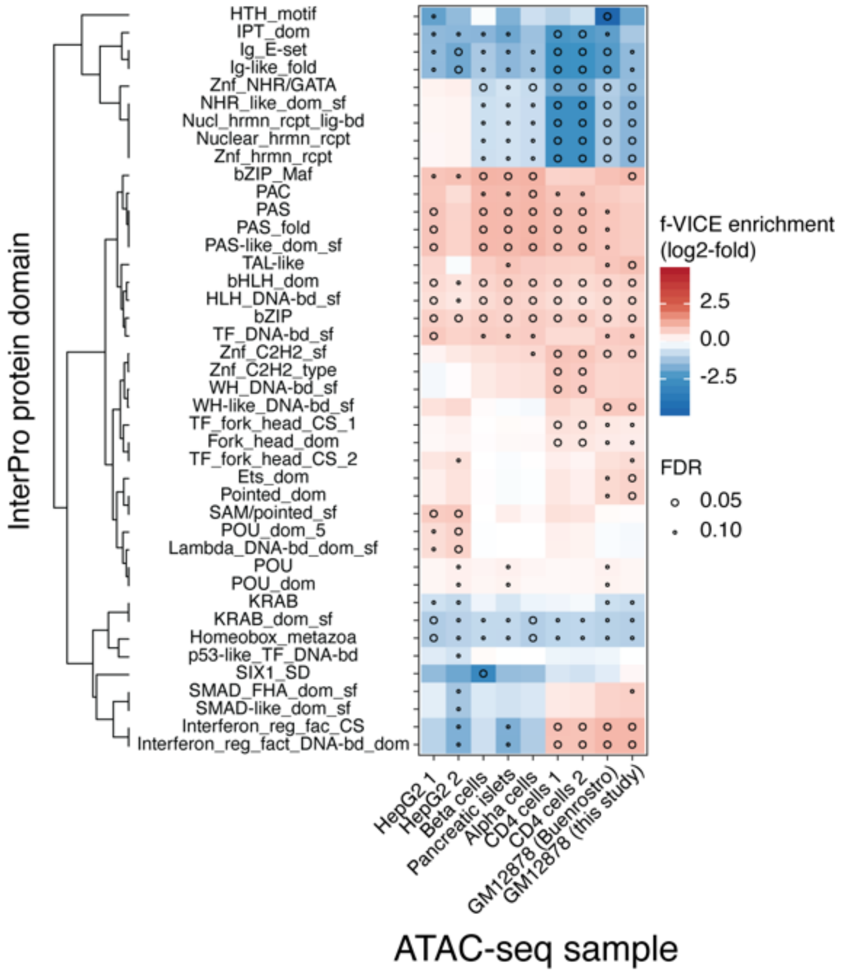


**Fig. S18. InterPro protein domain enrichments.**
InterPro protein domains f-VICE enrichments across samples.

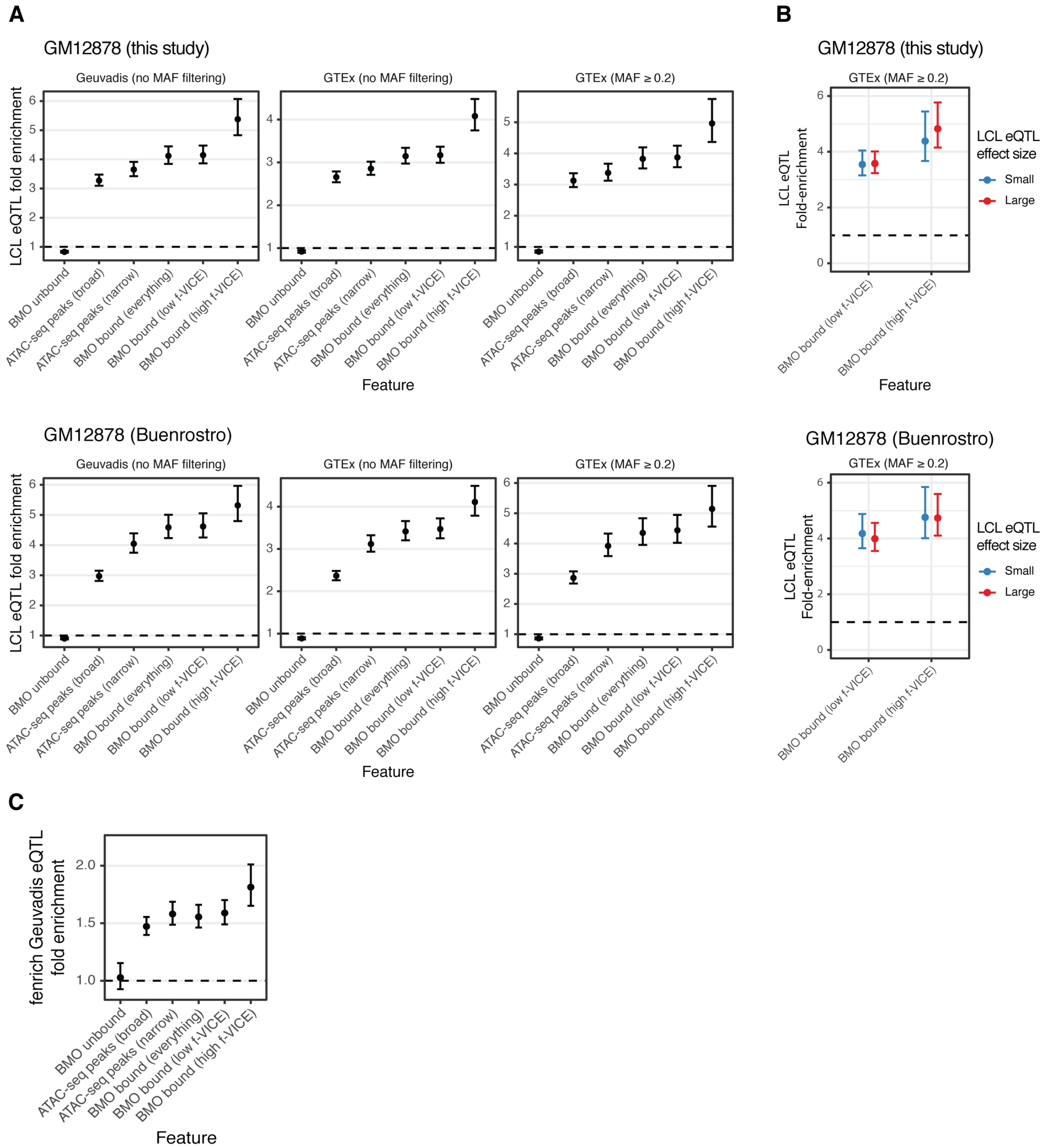


**Fig. S19**. **Enrichment of high and low f-VICE motifs in cis-eQTLs.**
(**A**) eQTL enrichments of different features across GM12878 ATAC-seq and lymphoblastoid cell lines (LCL) eQTL datasets. Enrichments are shown for GTEx with and without minor allele frequency (MAF) filtering to demonstrate that the observed results are not due to disproportionate representation of low MAF variants in any feature. (**B**) Enrichments of high and low f-VICE BMO predictions on high and low effect size GTEx eQTLs (above and below the median, respectively) across the two GM12878 datasets. (**C**) Geuvadis LCL eQTL enrichment calculated using QTL tools fenrich in our GM12878 dataset. Error bars in all plots represent the standard deviation of the effect size.

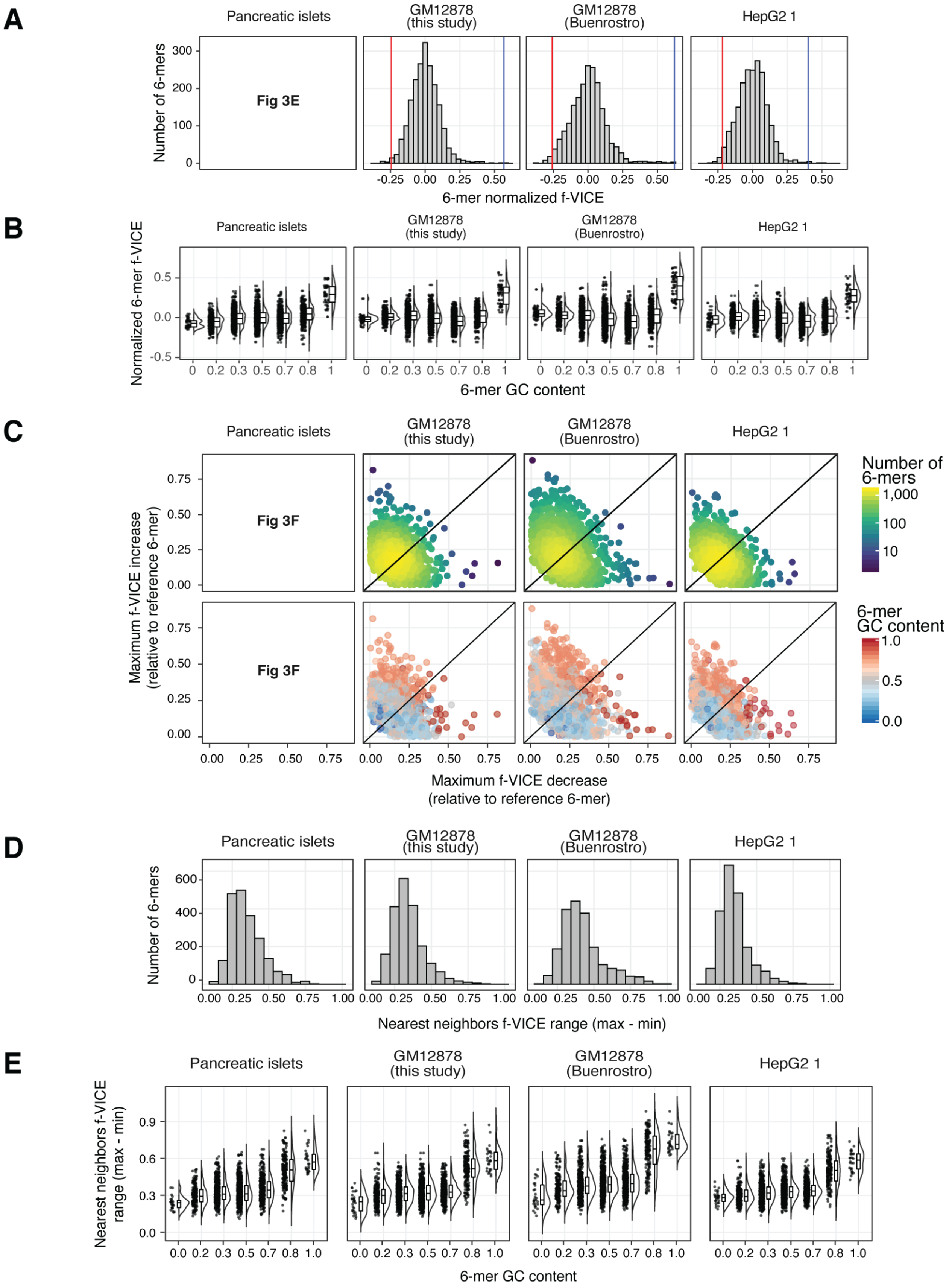

A
Pancreatic islets
GM12878 (this study)
GM12878 (Buenrostro)
HepG2 1
Number of 6-mers
Fig 3E
6-mer normalized f-VICE
B
Normalized 6-mer f-VICE
6-mer GC content
C
Maximum f-VICE increase (relative to reference 6-mer)
Fig 3F
Maximum f-VICE decrease (relative to reference 6-mer)
Number of 6-mers
1,000
100
10
6-mer GC content
D
Number of 6-mers
Nearest neighbors f-VICE range (max - min)
E
Nearest neighbors f-VICE range (max - min)
6-mer GC content

**Fig. S20. DNA 6-mers f-VICE analyses.**

(**A**) DNA 6-mers normalized f-VICE distributions across datasets. Horizontal lines represent the normalized f-IVCEs of the two 6-mers shown in Fig. 3E (CGCCCC in blue and CGACCC in red). (**B**) 6-mer f-VICEs as a function of GC content. (**C**) Scatter plot of f-VICE differences for all 6-mers relative to 1bp neighbors in sequence space (*i.e.* 6-mers with a Hamming distance of 1). (**D**) Distribution of the f-VICE range of each 6-mer relative to its 1bp neighbors in sequence space. (**E**) Distribution of f-VICE range as a function of GC content. Note that high GC content 6-mers are more likely to have immediate neighbors in sequence space with lower f-VICEs.

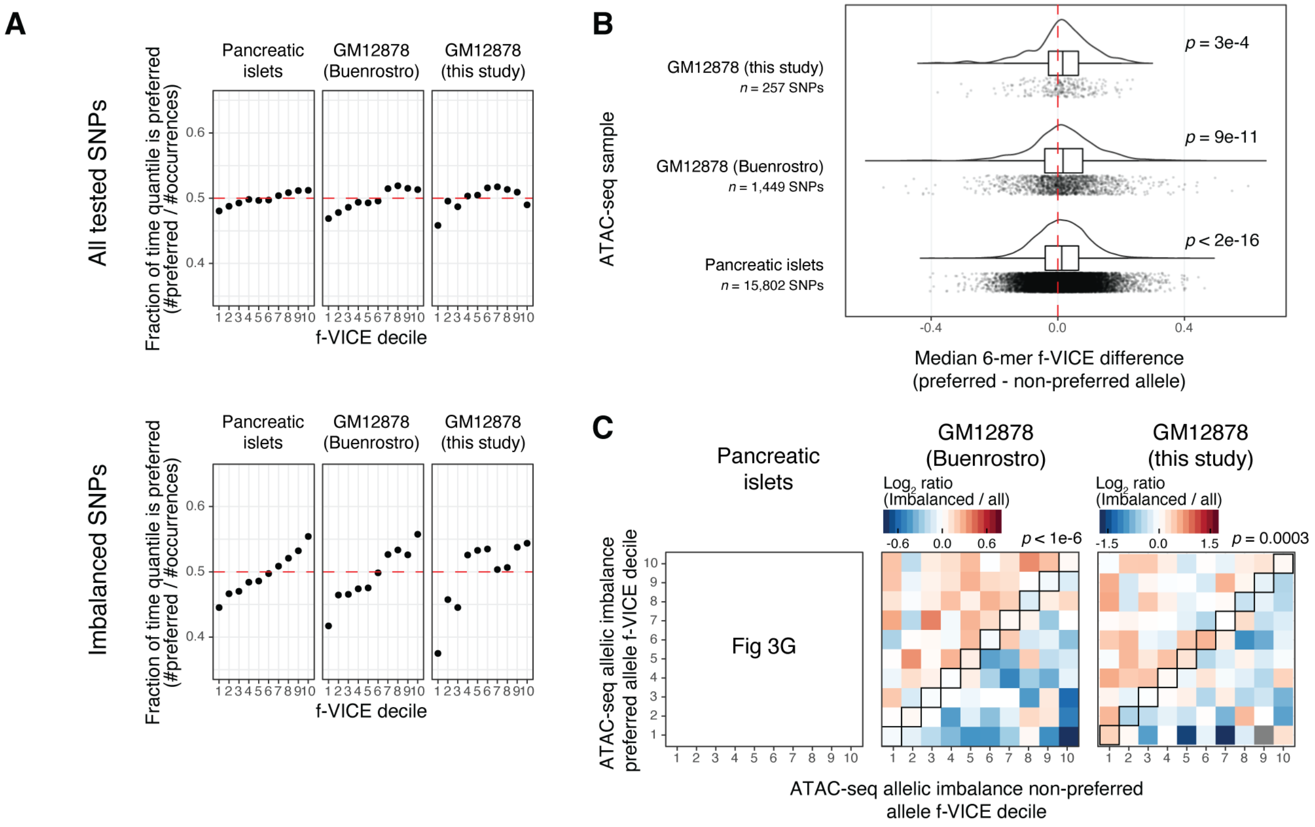


**Fig. S21. f-VICE allelic imbalance analyses.**
(**A**) Proportion of time the preferred ATAC-seq allele forms a 6-mer belonging to each f-VICE decile in all tested SNPs (upper) and all SNPs with significant allelic imbalance (lower). (**B**) Distribution of 6-mer f-VICE difference between the preferred and non-preferred alleles at loci with significant ATAC-seq imbalance. Each point corresponds to a DNA 6-mer overlapping a locus with allelic imbalance. Red dashed line corresponds to the expectation. P-values obtained from binomial tests. (**C**) f-VICE decile transition matrices. Each square corresponds to the ratio of imbalanced versus all tested SNPs. P-values obtained from permutation tests.

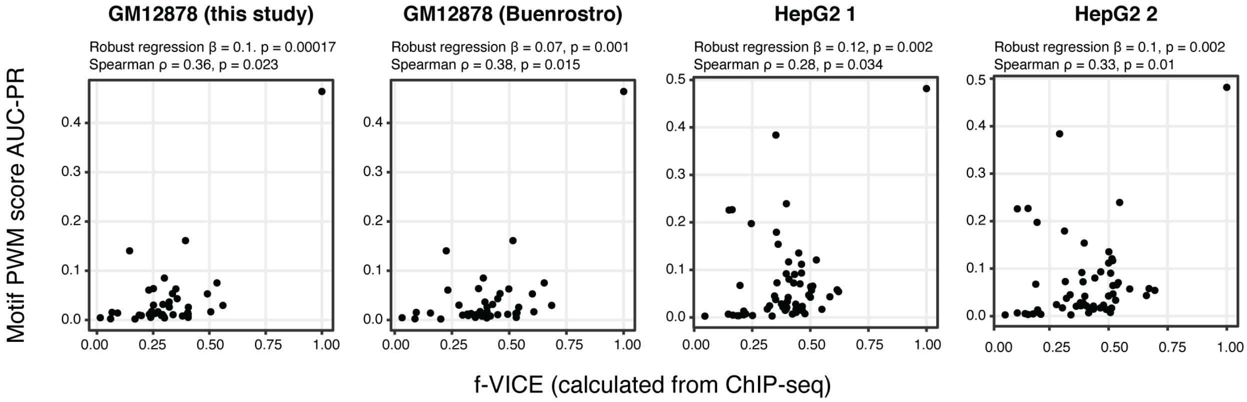


**Fig. S22. f-VICE and PWM score AUC-PR.**

Scatter plots of f-VICE and FIMO position weight matrix (PWM) score AUC-PR relative to ChIP-seq data. Robust linear regressions calculated using formula AUC-PR ~ f-VICE.

**Table S4.**
**FRAP recovery times from literature**

| Factor | Organism | Motif | FRAP recovery (s) | Reference |
|---|---|---|---|---|
| AHR | *Homo sapiens* | AHR_1 | 38 | (*69*) |
| AP1 | *Homo sapiens* | MA0476.1 | 600 | (*70*) |
| ARNT | *Homo sapiens* | ARNT_2 | 41 | (*69*) |
| CEBP | *Homo sapiens* | CEBPB_known5 | 32 | (*69*) |
| CREB | *Homo sapiens* | CREB3_1 | 100 | (*71*) |
| CTCF | *Homo sapiens* | CTCF_known2 | 660 | (*72*) |
| FOXA1 | *Mus musculus* | FOXA_known4 | 300 | (*73*) |
| MYC | *Homo sapiens* | MYC_known13 | 37 | (*69*) |
| NFKB | *Homo sapiens* | NFKB_known5 | 30 | (*74*) |
| NR3C1 | *Cercopithecus aethiops* | NR3C1_known18 | 30 | (*75*) |
| NR3C2 | *Homo sapiens* | NR3C2_1 | 30 | (*76*) |
| TP53 | *Homo sapiens* | TP53_4 | 20 | (*77*) |
| XBP | *Homo sapiens* | XBP1_2 | 30 | (*69*) |
